# Movement refinement and cholinergic regulation of outcome-dependent updating during skilled reaching

**DOI:** 10.64898/2026.10.29.762940

**Authors:** Nai-Hsing Yeh, Lily Prokop, John J Marshall, Anis Contractor

**Affiliations:** Department of Neuroscience, Northwestern University Chicago, IL 60613; Department of Psychiatry and Behavioral Sciences Feinberg School of Medicine, Northwestern University Chicago, IL 60613; Department of Neurobiology, Northwestern University Chicago, IL 60613

## Abstract

Motor skill learning transforms initially variable actions into precise and reproducible movements. Because conventional measures of task success obscure the distinct processes through which this refinement occurs, we combined detailed outcome scoring and kinematic analysis of single-pellet reaching in mice to resolve how individual components of the action change with repetition. Decomposing reaching into successive stages revealed improvements in targeting and retrieval, while grasp success-rate after pellet contact did not change detectably. Paw trajectories became more consistent across both successful and unsuccessful reaches, whereas changes in the mean speed profile were only detected in successful reaches. To examine how striatal cholinergic signaling contributes to movement refinement and the influence of recent outcomes on subsequent actions, we monitored acetylcholine (ACh) in vivo and inhibited cholinergic interneurons (CINs) during training. ACh was elevated before reach onset and declined during execution across all reach outcomes types. CIN inhibition had little effect on movement refinement across days but altered how recent outcomes influenced the next reach. In control animals, a failed attempt moved the reach away from the animal’s practiced successful trajectory but this did not occur after CIN inhibition, most clearly when the pellet had been grasped and then lost. These findings distinguish refinement of the practiced movement from outcome-dependent adjustments to its execution. Together, they raise the possibility that the pre-reach ACh elevation plays a role in enabling the evaluation of recent outcomes to update the execution of the next reach.

## Introduction

The goal of motor learning is to maximize an action’s desired outcome by refining the movements used to perform it. This iterative process involves dissociable changes in the individual components of the complex movement. For instance, in reaching for an object, successful performance requires accurate targeting, grasping, and retrieval, each of which can improve independently. While outcome alone cannot reveal which components of the action have improved or whether the underlying movement has become more consistent, the outcome of one trial may influence how the movement is performed on subsequent trials. Examining the individual stages of an action, its kinematics, and the influence of preceding outcomes can therefore help distinguish changes in motor execution from changes driven by recent experience.

The acquisition of a motor skill involves refinement of several features of movement that need not develop together. The consistency of a movement trajectory, the timing of its components, and the precision of fine movements can change over different timescales and depend on distinct contributions from cortical, striatal, and cerebellar circuits (Becker and Person, 2019; Lemke et al., 2019; Bova et al., 2021). Reach-to-grasp behavior exemplifies these distinctions: gross forelimb trajectories can stabilize while digit movements continue to improve, and movement consistency can increase even when overall retrieval success changes little (Bova et al., 2021). Thus, task outcome, a primary metric of motor skill learning, only captures part of the learning that occurs with practice. Distinguishing these aspects of motor learning is essential for understanding how the underlying neural circuits contribute to the acquisition and performance of skilled actions.

The dorsal striatum is well positioned to link the outcome of an action to changes in how that action is performed. Convergent cortical inputs provide information about movement and its context, while plasticity in corticostriatal circuits can modify the patterns of activity recruited through practice. Learning is accompanied by changes in corticostriatal activity associated with the refinement of task-relevant behavioral variability and by selective strengthening of cortical inputs from neurons recruited during training (Costa et al., 2004; Santos et al., 2015; Hwang et al., 2022). Dorsolateral striatal activity is required for the detailed kinematics of practiced movements, and action-specific striatal ensembles can influence the force of an ongoing forelimb action (Dhawale et al., 2021; Rodrigues-Vaz et al., 2025). Although some learned movements can be executed without motor cortex after training, their dependence on striatal circuitry persists, indicating that the mechanisms supporting acquisition and practiced performance are not identical (Kawai et al., 2015; Dhawale et al., 2021). Understanding striatal contributions to skill learning therefore requires distinguishing changes in movement execution from a narrower definition of response outcome. This involves determining how local signals regulate the processing of cortical input as movements are learned and performed.

Cholinergic signaling regulates how cortical and thalamic inputs engage striatal circuits. Acetylcholine (ACh) released by local cholinergic interneurons (CINs) modulates corticostriatal transmission and synaptic plasticity, while synchronized CIN activity can recruit dopamine release through nicotinic receptors on dopaminergic axons (Ding et al., 2010; Threlfell et al., 2012). Coordinated CIN activity accompanies transitions in movement, and direct measurements of ACh release reveal ongoing fluctuations as well as responses to salient events (Howe et al., 2019; Krok et al., 2023). The timing of cholinergic signaling relative to afferent activity is consequently an important consideration in understanding how the striatum processes information during behavior.

Here, we decomposed skilled reaching into its component actions and combined this analysis with quantitative tracking of paw movements, imaging of striatal ACh, and chemogenetic inhibition of CINs. Assessing targeting, grasping, and retrieval conditional on completion of the preceding movement stage revealed incremental improvements that evolved independently of success in the primary outcome measure. Training improved targeting and retrieval, while grasp success after pellet contact did not change detectably. Paw trajectories became more consistent across successful and unsuccessful outcomes, whereas changes in the speed profile were most evident in successful reaches. Striatal ACh was elevated before reach onset and declined during execution, in both early and late training. CIN inhibition with DREADDs (Designer Receptors Exclusively Activated by Designer Drugs) did not detectably impair the principal measures of movement refinement but altered how recent outcomes influenced the following reach, most clearly after a failed retrieval. These findings reveal distinct changes in movement with practice and suggest that striatal cholinergic signaling contributes to how recent outcomes influence subsequent actions.

## Results

To examine how the execution and outcome of skilled reaching change with practice, we trained mice on a single-pellet reaching task (**Figure 1A**). Following food restriction and two days of habituation, mice underwent two days of pre-training in which pellets were positioned progressively farther from the reaching aperture to establish reaching with the preferred forelimb. Mice then completed ten, daily training sessions, reaching through a narrow aperture to grasp a pellet and retrieve it to the mouth. Each session ended after 20 pellets were successfully retrieved or 10 minutes had elapsed. Reaching was recorded with high-speed video, allowing individual attempts to be scored by outcome and paw movements to be tracked using DeepLabCut. We compared early training (days 1–2) with late training (days 9–10) to assess changes in both task performance and movement execution.

**Figure 1.**
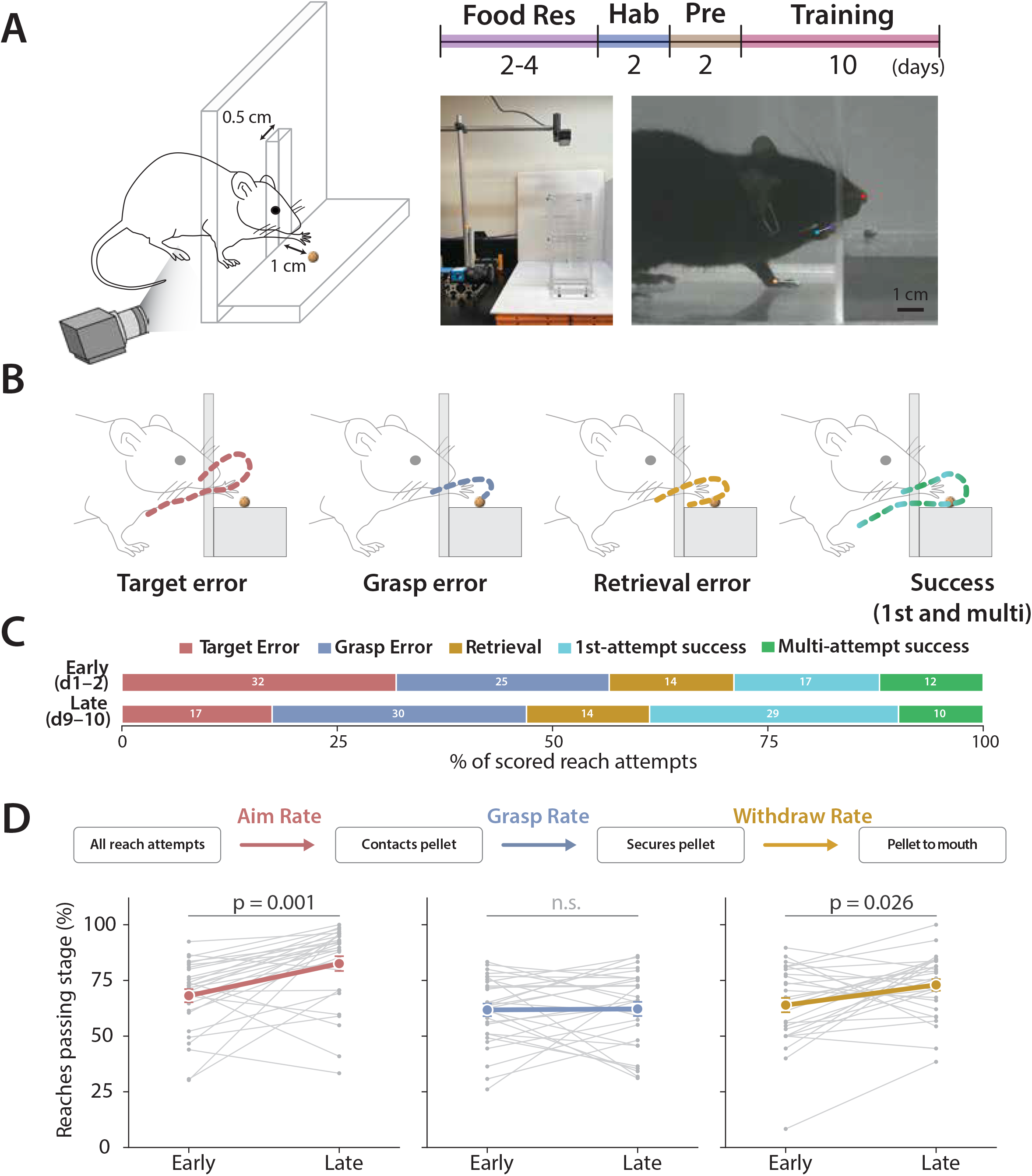
Training selectively improves targeting and retrieval during skilled reaching. **(A)** Single-pellet reaching apparatus and training timeline. Mice reached through a 0.5-cm-wide aperture for a pellet positioned 1 cm from the front wall. Following 2–4 days of food restriction, mice underwent two days of habituation, two days of pre-training, and ten days of training. Pictures show the apparatus and a representative high-speed video frame with tracked landmarks. **(B)** Schematic of reach outcomes. Target errors: failure to contact the pellet; grasp errors: pellet contact without a secure grasp, including displacement of the pellet; retrieval errors: pellet secured but lost before consumption; successes: pellet returned to the mouth and consumed. Successful retrievals were further classified as first-attempt or multiple-attempt successes. Dashed lines illustrate representative paw trajectories. **(C)** Distribution of reach outcomes during early (days 1–2) and late (days 9–10) training. Stacked bars show the mean percentage of attempts in each category, calculated within each mouse; labels give rounded percentages. Early versus late comparisons: target errors, W = 61.0, P_Holm = 0.0018; grasp errors, W = 128.0, P_Holm = 0.160; retrieval errors, W = 209.0, P_Holm = 0.865; first-attempt successes, W = 84.0, P_Holm = 0.012; multiple-attempt successes, W = 141.0, P_Holm = 0.202. Individual-animal values are shown in Figure S1A. (**D**) Conditional performance at successive stages of reaching. Top, schematic defining aim rate as the fraction of all attempts contacting the pellet, grasp rate as the fraction of pellet-contacting attempts securing it and withdraw rate as the fraction of successful grasps returning the pellet to the mouth. Bottom, early and late rates for individual mice. Grey lines connect paired observations; colored symbols and lines show mean ± s.e.m. Aim rate increased from 68.2 ± 2.9% to 82.6 ± 3.3% (W = 61.0, P = 3.7 × 10⁻⁴, P_Holm = 0.0011); grasp rate was 61.8 ± 2.8% and 62.2 ± 3.2% (W = 204.0, P = 0.78, P_Holm = 0.78); withdraw rate increased from 63.9 ± 3.2% to 73.0 ± 2.6% (W = 104.0, P = 0.013, P_Holm = 0.026). Plot annotations show Holm-adjusted P values; n.s., not significant. Data comprise 5,215 scored reach attempts from 29 wild-type mice, including 2,182 early and 3,033 late attempts. The mouse was the unit of analysis throughout. Comparisons used two-sided paired Wilcoxon signed-rank tests, with Holm–Bonferroni correction applied separately to the five outcome categories in (**C**) and the three conditional rates in (**D**).

Across 29 mice, successful pellet retrieval increased from 28.9 ± 2.6% of reach attempts during early training to 38.7 ± 3.2% during late training (P = 0.0065, Wilcoxon signed-rank test). However, improvement varied substantially between individual animals: 18 of 29 mice (62%) increased their success rate, and among those, only 11 improved by more than 10 percentage points.

However, success rate alone does not indicate which components of the action changed, or whether mice whose success did not increase had nonetheless refined their reaching. We therefore classified each of the 5,215 reach attempts into five mutually exclusive outcome categories based on the stage at which the action succeeded or failed: target errors (failure to contact the pellet), grasp errors (contact without successful grasp, including attempts in which the pellet was displaced), retrieval errors (pellet grasped but dropped before consumption), multiple-attempt successes (pellet obtained after more than one reach), and first-attempt successes (pellet obtained on the initial reach; **Figure 1B**).

We first asked how the distribution of reach outcomes changed between early and late training (**Figure 1C**). For each mouse, we calculated the fraction of reaches assigned to each of the five outcome categories. Percentages were calculated within individual animals because the number of attempts varied between mice, ensuring that animals making more reaches did not contribute disproportionately to the comparison. Training shifted the distribution toward fewer target errors and more first-attempt successes. Target errors decreased from 31.8 ± 2.9% during early training to 17.4 ± 3.3% during late training, whereas first-attempt successes increased from 16.9 ± 2.1% to 28.9 ± 3.6% (P = 0.0018 and P = 0.012, respectively; Wilcoxon signed-rank tests with Holm correction). The proportions of retrieval errors and multiple-attempt successes did not change significantly (P = 0.865 and P = 0.202, respectively). Grasp errors increased numerically from 24.7 ± 1.5% to 29.5 ± 2.3%, although this difference was not significant after correction for multiple comparisons (P = 0.160). Individual-animal values for all five outcome categories are shown in **Figure S1A**.

Interpreting these changes requires accounting for the sequential structure of the reaching task. While outcome categories marked the stage at which an attempt ended, the progression to each stage depended on successful completion of the preceding one. Thus, a grasp error can occur only after the paw contacts the pellet, and a retrieval error only after the pellet has been grasped. The fraction of all attempts ending in a particular error therefore depends both on performance at that stage and on how many attempts reach it. As targeting improves, more reaches contact the pellet, creating more opportunities for both successful grasps and grasp errors. Consequently, grasp errors can account for a larger proportion of all attempts even when the probability of grasping the pellet after contact is unchanged. Equally, an improvement at a later stage can be masked when more attempts reach it: better targeting and grasping send more pellets to the retrieval stage, so the number of retrieval errors can stay constant even as retrieval itself improves.

To distinguish these effects, we analyzed the same attempts as a nested cascade, evaluating each stage only on the attempts that completed the preceding one (**Figure 1D**). For each animal and training phase, we calculated three conditional probabilities: the aim rate, defined as the fraction of all attempts that contacted the pellet; the grasp rate, defined as the fraction of pellet contacts that resulted in a successful grasp; and the withdraw rate, defined as the fraction of successful grasps in which the pellet was returned to the mouth. This analysis separated performance at each component of the action from changes in the number of attempts progressing to it. The cascade also provides an exact decomposition of overall task success: within each animal and training phase, the product of the aim, grasp, and withdraw rates equals the overall probability of successful pellet retrieval.

This analysis revealed a clear stage-specific pattern of learning (**Figure 1D**). Aim performance improved from 68.2 ± 2.9% during early training to 82.6 ± 3.3% during late training, with 24 of 29 animals showing an increase (P = 0.0011, Wilcoxon signed-rank test with Holm correction). Withdrawal performance also improved, from 63.9 ± 3.2% to 73.0 ± 2.6%, with 20 of 29 animals improving (P = 0.026). In contrast, grasp performance did not change detectably (61.8 ± 2.8% versus 62.2 ± 3.2%; P = 0.78), with approximately equal numbers of animals showing increases and decreases (16 of 29 improved). The 95% confidence interval for the paired change in grasp performance ranged from −6.7 to +7.5 percentage points, excluding mean improvements as large as those observed in targeting and withdrawal. Thus, the numerical increase in grasp errors among all attempts did not indicate poorer grasp performance after contact. Improved targeting allowed more attempts to progress to the grasp stage, increasing the opportunities for grasp errors even though conditional grasp success remained similar.

To quantify how changes at successive stages contributed to the observed outcome distribution, we decomposed the training-related change in the proportion of trials ending with a given error type into changes in performance at preceding stages and performance at the error stage itself (**Figure S1B**). For grasp errors, improved targeting contributed an increase of 5.3 ± 1.7 percentage points (P = 0.002, Wilcoxon signed-rank test), whereas the contribution from changes in grasp performance was −0.5 ± 2.4 percentage points (P = 0.85). Together, these accounted for the observed 4.8 percentage-point increase in grasp errors among all attempts. Retrieval errors illustrated how the same sequential structure could conceal an improvement. Their overall proportion changed little with training (−0.2 ± 2.1 percentage points; P = 0.865), despite improved retrieval in the conditional analysis. Increased progression through the preceding stages contributed +3.1 ± 1.4 percentage points (P = 0.053), while improved retrieval contributed −3.3 ± 1.5 percentage points (P = 0.031). That is, animals were more likely to make a successful retrieval once they grasped the pellet, however this improvement was not apparent when measuring the overall proportion of trials ending in a retrieval error. The opposing effects of increased progression to the retrieval stage and improved withdrawal largely offset one another, masking the improvement in retrieval performance.

These results show that individual components of skilled reaching improve differently over ten days of training. Mice became more successful at directing the paw to the pellet and returning a secured pellet to the mouth, while the probability of grasping the pellet after contact did not change detectably. Overall success rate as a measure of performance does not resolve these differences, and expressing errors as a proportion of all attempts can obscure improvements or suggested deficits because the number of attempts reaching each stage changes with training. Having identified where performance improved within the action sequence, we next examined how practice altered the consistency and timing of the underlying paw movements.

Practice can increase movement consistency even when overall task success changes little (Bova et al., 2021). We therefore quantified trial-to-trial variability in paw trajectories separately for each of the five reach outcomes, comparing early (days 1–2) and late (days 9–10) training. Paw position was tracked in the sagittal plane to measure forward displacement relative to the front panel and height above the platform (**Figure 2A**). Individual trajectories were time-normalized across the outward reach and withdrawal and aligned to the point at which the paw crossed the front panel, providing a common physical landmark for comparison across trials and animals.

**Figure 2.**
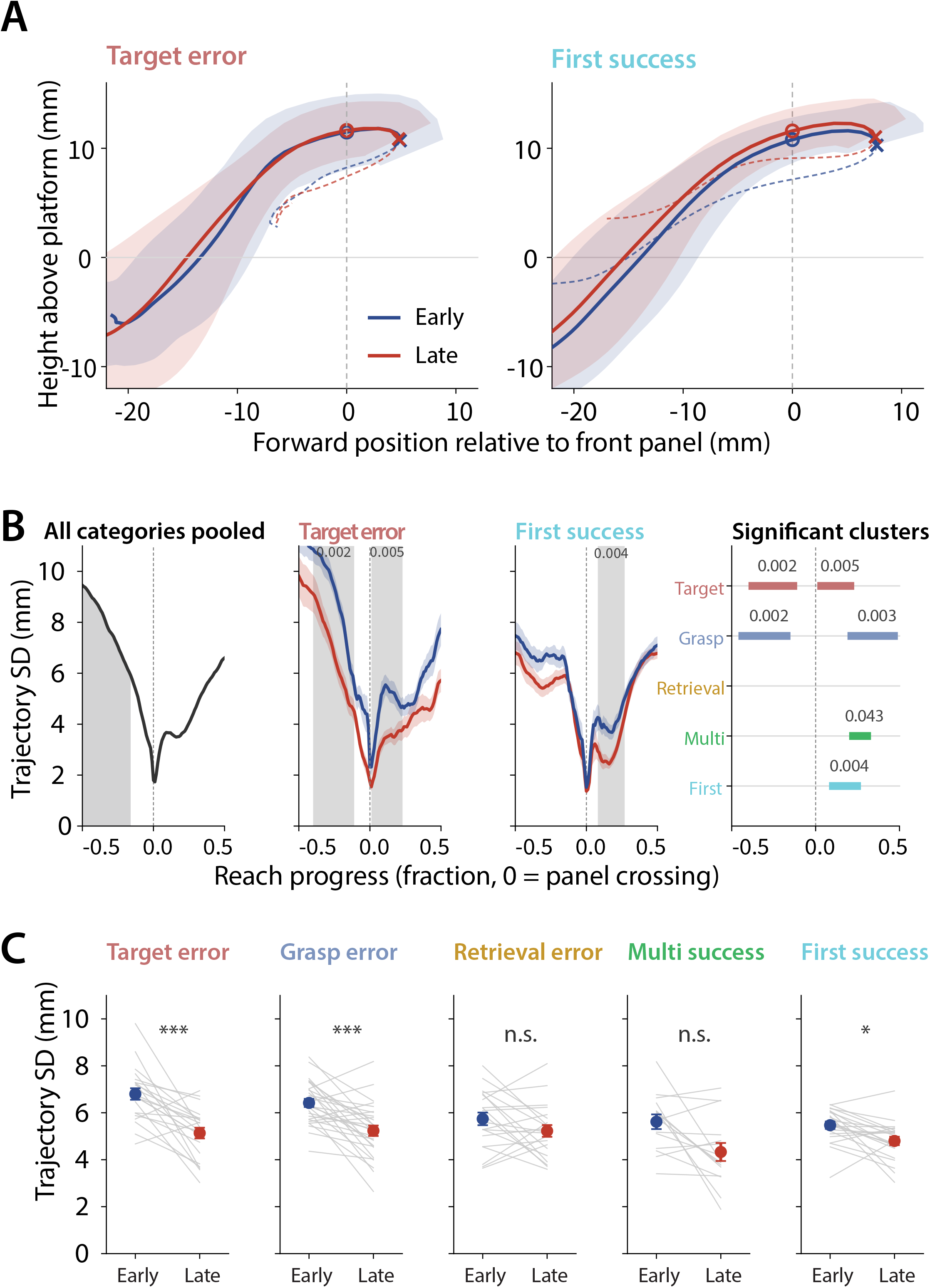
Practice reduces reach trajectory variability across successful and unsuccessful outcomes. **(A)** Mean paw trajectories in the sagittal plane for target errors and first-attempt successes during early (days 1–2, blue) and late (days 9–10, red) training. Forward position is measured relative to the front panel (x = 0), and height relative to the platform surface (y = 0). Both axes use the same spatial scale. Trajectories were time-normalized, aligned to front-panel crossing, and averaged within sessions, within mice, and then across mice. Open circles indicate panel crossing; crosses indicate maximum forward extent. Shading shows ±1 s.d. across mice perpendicular to the mean outward trajectory. Withdrawal is shown as a thin dashed line without shading. The initial approach before x = −22 mm is omitted for clarity. n = 22 mice for each outcome category. (**B**) Across-trial trajectory variability along the reach. Left, variability pooled across outcome categories and training phases, illustrating greater dispersion near reach onset and a minimum near panel crossing. Shading identifies the initial approach. Middle panels: variability for target errors and first-attempt successes during early and late training, shown as mean ± s.e.m. across mice. Grey bands indicate significant clusters from paired comparisons between training phases. Right, locations of significant clusters across all five outcome categories; colors identify outcomes, and grey lines indicate categories without a significant cluster. Cluster-mass permutation tests used pointwise paired t-statistics, a two-sided cluster-forming threshold of P < 0.05, and 5,000 sign-flips. Target errors showed significant clusters before and after crossing (−0.40 to −0.11, P = 0.0024; 0.01 to 0.23, P = 0.0052), as did grasp errors (−0.46 to −0.15, P = 0.0016; 0.19 to 0.49, P = 0.0034). Significant clusters occurred after crossing for first-attempt successes (0.08 to 0.27, P = 0.0038) and multiple-attempt successes (0.20 to 0.33, P = 0.043). No significant cluster was detected for retrieval errors. Intervals are expressed in normalized reach progress, with panel crossing at zero. (**C**) Across-trial trajectory variability averaged over the complete trajectory for each mouse during early and late training calculated as the mean of the across-trial positional standard deviations in the forward and vertical coordinates. Grey lines connect paired observations; colored symbols show mean ± s.e.m. Target errors, 6.80 to 5.14 mm (n = 22, W = 12, P_Holm = 1.7 × 10⁻⁴); grasp errors, 6.42 to 5.23 mm (n = 28, W = 46, P_Holm = 5.4 × 10⁻⁴); retrieval errors, 5.74 to 5.23 mm (n = 21, W = 78, P_Holm = 0.20); first-attempt successes, 5.47 to 4.80 mm (n = 22, W = 41, P_Holm = 0.012); multiple-attempt successes, 5.62 to 4.33 mm (n = 15, W = 21, P_Holm = 0.051). Comparisons used two-sided Wilcoxon signed-rank tests with Holm– Bonferroni correction across the five categories. Sessions containing fewer than three scored trials in an outcome category were excluded from variability estimates for that category. Each mouse contributed one value per training phase, and the mouse was the unit of analysis. *P < 0.05, **P < 0.01, ***P < 0.001; n.s., not significant. Significance symbols in (**C**) refer to Holm-adjusted P values.

Trajectory variability differed across the movement, decreasing as the paw approached the aperture and increasing again during withdrawal (**Figure 2B**). The minimum at panel crossing partly reflects alignment to this landmark. We therefore assessed training-related changes both across the complete trajectory and at successive points along the movement. Averaged over the whole reach, trajectory variability decreased with training in both unsuccessful and successful attempts (**Figure 2C**). Variability fell in target errors, from 6.80 to 5.14 mm (n = 22 mice; P_Holm = 1.7 × 10⁻⁴), and grasp errors, from 6.42 to 5.23 mm (n = 28; P_Holm = 5.4 × 10⁻⁴). First-attempt successes also became more consistent, with variability decreasing from 5.47 to 4.80 mm (n = 22; P_Holm = 0.012; Wilcoxon signed-rank tests with Holm correction across the five categories). Variability declined numerically in multiple-attempt successes, from 5.62 to 4.33 mm, but did not reach significance after correction (n = 15; P_Holm = 0.051). Retrieval errors showed no detectable change (5.74 to 5.23 mm; n = 21; P_Holm = 0.20).

Analysis along the trajectory identified phases of the reach where variability declined between early and late training (**Figure 2B**; cluster-mass permutation tests). For target and grasp errors, significant reductions occurred both before and after panel crossing (target errors: P = 0.0024 and P = 0.0052; grasp errors: P = 0.0016 and P = 0.0034). For successful reaches, significant reductions were detected only after crossing (first-attempt successes: P = 0.0038; multiple-attempt successes: P = 0.043). No significant interval was detected for retrieval errors. These results show that increased movement consistency is not restricted to reaches that successfully retrieve the pellet. Target errors, grasp errors, and first-attempt successes all became less variable with practice, demonstrating refinement of the movement even when an attempt remained unsuccessful. Multiple-attempt successes showed a localized reduction in variability after panel crossing, although the whole-reach comparison did not reach significance after correction.

Trajectory variability captures the spatial consistency of the reach but does not reveal how the movement is organized in time. We therefore compared instantaneous paw speed during early and late training, again aligning reaches to aperture crossing (**Figure 3A**). Significant changes in the speed profile were detected only in successful reaches. During first-attempt successes, paw speed decreased over a broad interval beginning shortly after aperture crossing and extending into withdrawal (0.10–0.43 s, P < 0.0001; cluster-mass permutation test). Multiple-attempt successes showed a reduction over a narrower interval (0.20–0.29 s, P = 0.023). No significant interval was detected for target, grasp, or retrieval errors. Thus, while trajectory consistency improved across several successful and unsuccessful outcomes, detectable changes in the speed profile only occurred in reaches that successfully retrieved the pellet.

**Figure 3.**
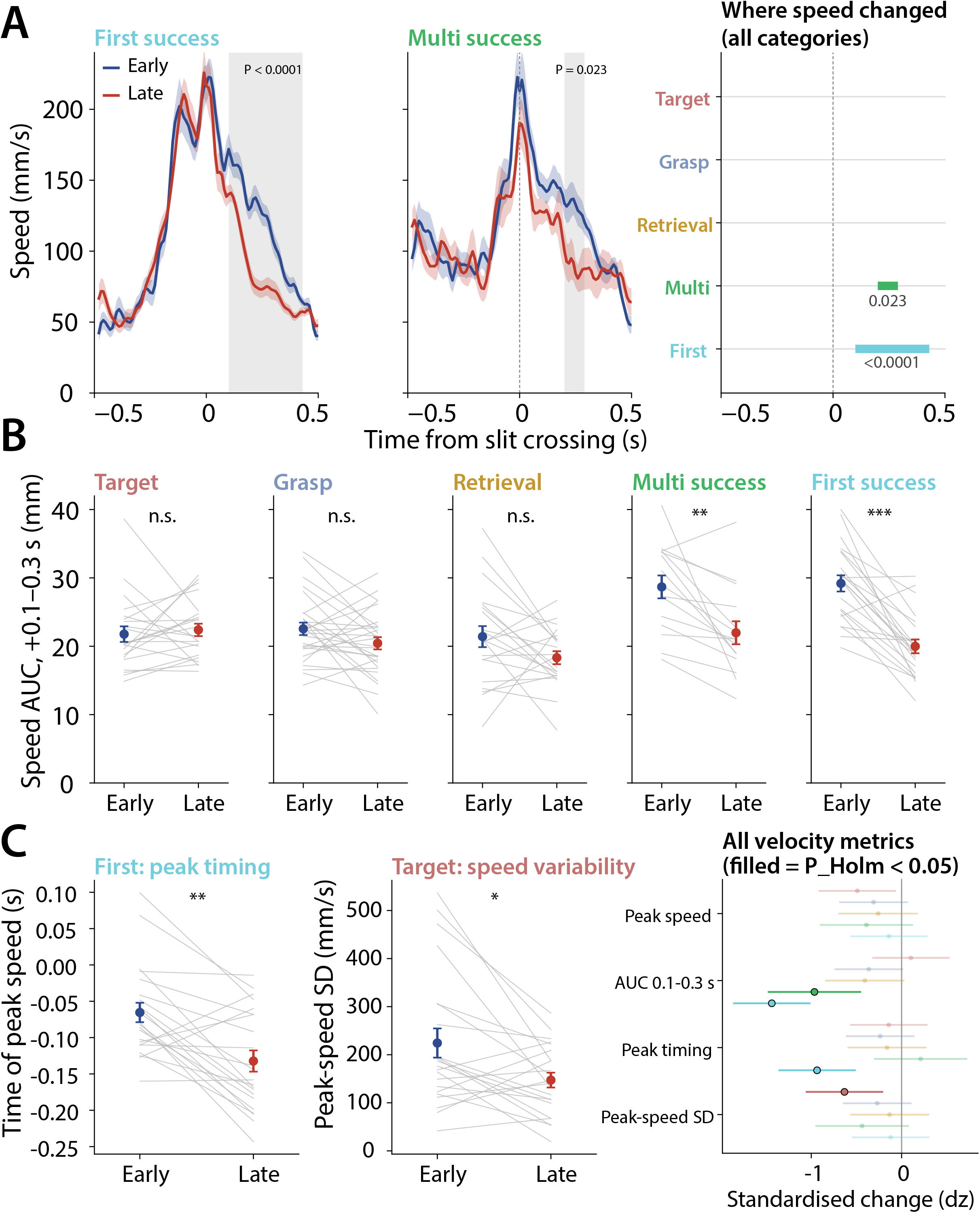
Training alters the speed profile of successful reaches and the consistency of peak speed in target errors. **(A)** Paw speed during early (days 1–2, blue) and late (days 9–10, red) training, aligned to aperture crossing (t = 0). Traces show mean ± s.e.m. across mice for first-attempt successes (n = 22) and multiple-attempt successes (n = 15). Grey bands indicate significant clusters from paired comparisons between training phases using pointwise t-statistics and cluster-mass permutation tests with 5,000 sign-flips. Paw speed decreased over +0.10 to +0.43 s in first-attempt successes (P < 0.0001) and +0.20 to +0.29 s in multiple-attempt successes (P = 0.023). Right, locations of significant clusters across outcome categories. No significant cluster was detected for target, grasp, or retrieval errors. **(B)** Integral of paw speed over +0.1 to +0.3 s after aperture crossing, expressed as distance traveled during that interval. Values were calculated per mouse and training phase. First-attempt successes decreased from 29.2 ± 1.2 to 20.0 ± 1.0 mm (n = 22, W = 6, P_Holm = 3.0 × 10⁻⁵), and multiple-attempt successes decreased from 28.7 ± 1.7 to 22.0 ± 1.7 mm (n = 15, W = 6, P_Holm = 0.0034). No significant change was detected in target errors (P_Holm = 0.34), grasp errors (P_Holm = 0.26), or retrieval errors (P_Holm = 0.26). Comparisons used two-sided Wilcoxon signed-rank tests with Holm–Bonferroni correction across the five outcome categories. **(C)** Left, timing of peak paw speed relative to aperture crossing in first-attempt successes. Peak speed occurred earlier with training, shifting from −65 ± 13 to −132 ± 15 ms (n = 22, W = 25, P_Holm = 0.0021). Middle, across-trial standard deviation of peak speed in target errors, which decreased from 224 ± 30 to 147 ± 15 mm/s (n = 22, W = 47, P_Holm = 0.042). Right, standardized early-to-late changes in four speed metrics across all five outcome categories, shown as Cohen’s dz with 95% confidence intervals. Filled symbols indicate significant changes after Holm correction. Mean peak speed and peak deceleration did not change significantly in any category (all P_Holm ≥ 0.14 and ≥ 0.30, respectively). Sessions containing fewer than three scored trials in an outcome category were excluded from analyses of that category. Each mouse contributed one value per training phase, and the mouse was the unit of analysis. Summary values are mean ± s.e.m.

To quantify the reduction in speed after aperture crossing, we integrated paw speed over the +0.1 to +0.3 s interval, which was defined from the interval identified by the cluster-based analysis above (**Figure 3B**). The integral of paw speed provides a measure of the distance traveled during this interval. In first-attempt successes, this distance decreased by 32%, from 29.2 ± 1.2 to 20.0 ± 1.0 mm (n = 22 mice; W = 6, P_Holm = 3.0 × 10⁻⁵; Wilcoxon signed-rank test). Multiple-attempt successes showed a 23% reduction, from 28.7 ± 1.7 to 22.0 ± 1.7 mm (n = 15; W = 6, P_Holm = 0.0034). No detectable change occurred in target errors (P_Holm = 0.34), grasp errors (P_Holm = 0.26), or retrieval errors (P_Holm = 0.26).

We next examined whether training changed the magnitude, timing, or trial-to-trial consistency of peak paw speed (**Figure 3C**). In first-attempt successes, peak speed occurred 67 ms earlier relative to aperture crossing, shifting from −65 ± 13 ms during early training to −132 ± 15 ms during late training (n = 22; W = 25, P_Holm = 0.0021). Successful first attempt reaches therefore showed a distinct speed profile with maximum speed attained early in the reach. Target errors showed a different change, with the trial-to-trial standard deviation of peak speed decreasing by 34%, from 224 ± 30 to 147 ± 15 mm/s (n = 22; W = 47, P_Holm = 0.042). Mean peak speed did not change significantly in any outcome category after correction for multiple comparisons (all P_Holm ≥ 0.14), nor did peak deceleration (all P_Holm ≥ 0.30).

These analyses show that practice changes both the consistency of paw trajectories and the timing of movement. In first-attempt successes, the paw attained peak speed earlier without a detectable change in its magnitude, followed by lower speed after aperture crossing. Target errors also became more consistent in their peak speed, despite no detectable change in the mean speed profile. These changes complement the reductions in trajectory variability, showing that practice involves the refinement of spatial reproducibility, temporal organization, and trial-to-trial consistency of the movement, each measurable independently of changes in the overall success rate.

The changes in movement consistency and timing raise the question of how striatal signals are organized during the acquisition and performance of skilled reaching, and whether these signals change in parallel to refinements in movement variability and dynamics. ACh regulates corticostriatal transmission and plasticity (Abudukeyoumu et al., 2019), and therefore can potentially modulate inputs recruited during an action and changes in striatal processing with experience. To examine this relationship, we first measured the time course of ACh fluctuations around individual reaches and asked whether it differed with reach outcome or training stage. We expressed the genetically encoded sensor GRAB-rACh3.0 in dorsolateral striatum and imaged bulk fluorescence through a gradient-index lens during single-pellet reaching, yielding 700 scored reaches across 35 sessions from six mice (**Figure 4A & B**).

**Figure 4.**
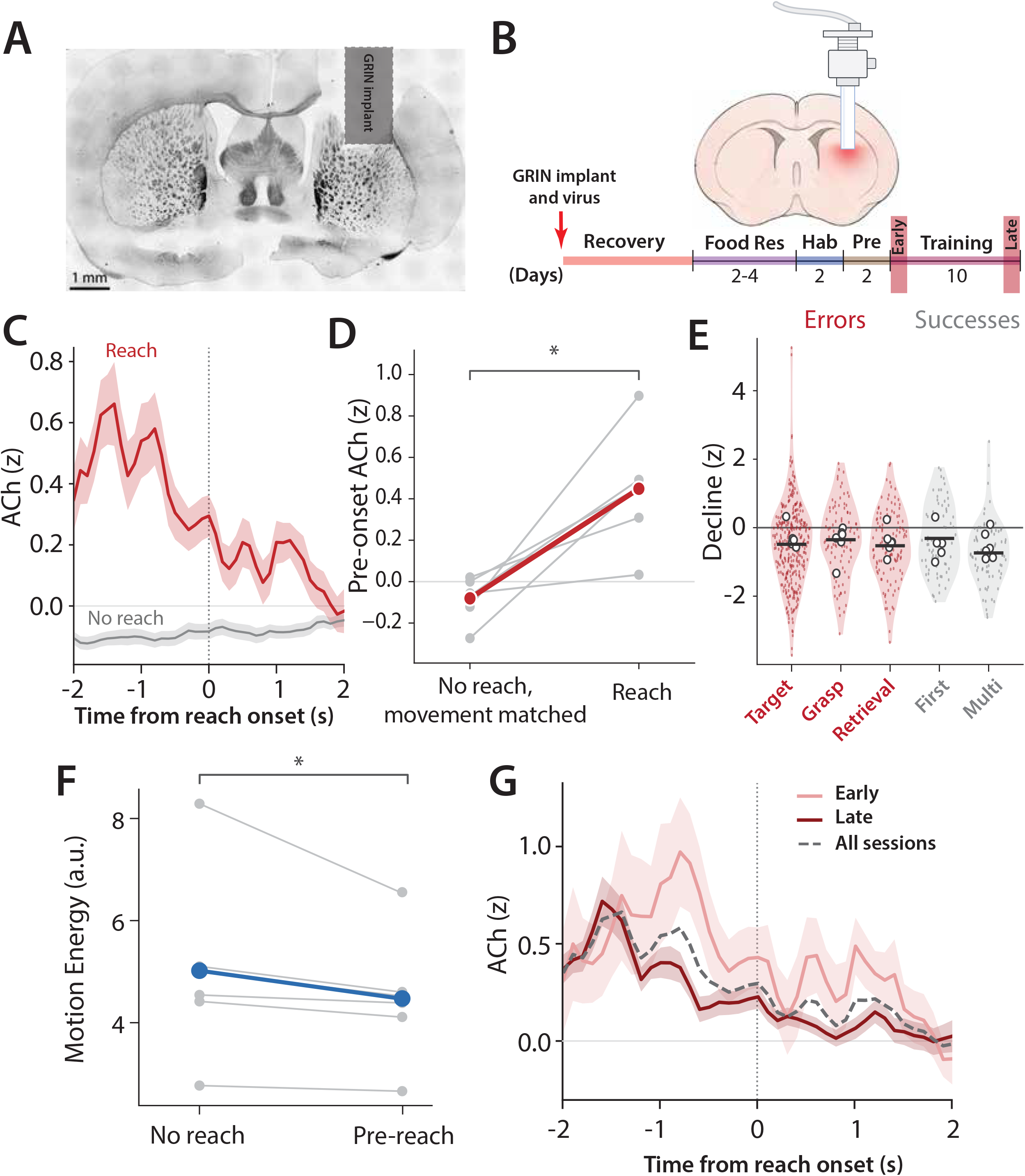
Striatal acetylcholine is elevated before reaching and declines during execution. (A) Representative coronal brain section showing the approximate location of the implanted GRIN lens in the dorsolateral striatum. The shaded region indicates the approximate lens placement. (**B**) Experimental timeline. A virus-coated GRIN lens was implanted in dorsolateral striatum to deliver GCaMP8f and GRAB-rACh3.0 at the imaging site. Following recovery, mice underwent food restriction, habituation, pre-training, and ten days of single-pellet reaching training. Analyses compare early (days 1–2) and late (days 9–10) training. (**C**) GRAB-rACh3.0 fluorescence aligned to reach onset (t = 0, first forward paw movement). Traces show mean ± s.e.m. across sessions after whole-session z-scoring of ΔF/F. Red, scored reaches; grey, reach-free comparison windows sampled from the same recordings at least 4 s from any scored reach, with 20 samples per reach. Pre-onset fluorescence was measured over −2.0 to −0.6 s; the decline was defined as mean fluorescence over +1.0 to +2.0 s minus the pre-onset value. Across 35 sessions, pre-onset fluorescence was 0.507 ± 0.097 z for reaches and −0.108 ± 0.023 z for reach-free windows (W = 32, P = 1.6 × 10⁻⁷). The corresponding changes were −0.406 ± 0.094 and +0.041 ± 0.019 z (W = 79, P = 3.8 × 10⁻⁵). Mouse-level comparisons gave pre-onset values of 0.487 ± 0.148 versus −0.105 ± 0.029 z (W = 0, P = 0.031) and changes of −0.396 ± 0.167 versus +0.042 ± 0.024 z (W = 2, P = 0.094; n = 6 mice). Tests were two-sided Wilcoxon signed-rank comparisons. (**D**) Pre-onset fluorescence during reaching versus movement-matched reach-free windows. Pre-onset fluorescence was −0.085 ± 0.037 z for matched reach-free windows and +0.442 ± 0.115 z for reaches (W = 0, P = 0.031). (**E**) Change in fluorescence from pre-onset to execution, separated by reach outcome; red denotes unsuccessful attempts and grey denotes successful retrievals. Violins are kernel densities over the individual reaches, black horizontal ticks are means across reaches and open circles are the mice. Target errors, −0.485 ± 0.063 z (344 reaches); grasp errors, −0.351 ± 0.112 z (91); retrieval errors, −0.528 ± 0.095 z (115); first-attempt successes, −0.314 ± 0.114 z (75); multiple-attempt successes, −0.738 ± 0.107 z (75). Category sample sizes differ substantially, so the densities are not equally well estimated. Mouse-level comparisons detected no difference among categories (Friedman test, χ² = 3.04, P = 0.55; n = 5 mice with all categories represented) or between successful and unsuccessful reaches (Wilcoxon signed-rank test, W = 4, P = 0.219; n = 6). (**F**) Motion energy during pre-onset and reach-free windows, calculated from frame-to-frame changes in a body region. Motion energy was lower before reaching, with a mean within-animal pre-onset/reach-free ratio of 0.909 ± 0.026 (W = 0, P = 0.031; two-sided Wilcoxon signed-rank test; n = 6 mice). (**G**) Reach-aligned fluorescence during early training (light red; 12 sessions, 71 reaches) and late training (dark red; 23 sessions, 629 reaches), shown as mean ± s.e.m. across sessions. The dashed grey trace shows the all-session average from (B). Pre-onset fluorescence did not differ detectably between phases (0.640 ± 0.252 versus 0.437 ± 0.070 z; Mann–Whitney U = 154, P = 0.59), nor did the decline (−0.448 ± 0.189 versus −0.383 ± 0.107 z; U = 149, P = 0.72). The paired mouse-level comparison of the decline was also not significant (W = 9, P = 0.84; n = 6). Data comprise 700 scored reaches across 35 sessions from six mice. Session-level summaries and tests are distinguished from mouse-level analyses throughout. Summary values are mean ± s.e.m.

Aligning the ACh signal to reach onset revealed an elevation before the first forward paw movement, followed by a decline during execution (**Figure 4C**). The mean signal peaked approximately 1.4 s before reach onset. Because the elevation extended into the pre-onset period, using that period as a local baseline could obscure the signal. We therefore z-scored fluorescence over the whole session and compared reach-aligned signals with signals aligned to reach-free time points drawn from the same recording. Pre-onset ACh, measured from −2.0 to −0.6 s, was higher than this reach-free comparison in all six mice (+0.487 ± 0.148 versus −0.105 ± 0.029 z; W = 0, P = 0.031, Wilcoxon signed-rank test). The elevation was also evident in 32 of 35 sessions (+0.507 ± 0.097 versus −0.108 ± 0.023 z; W = 32, P = 1.6 × 10⁻⁷). The signal subsequently declined between the pre-onset window and +1.0 to +2.0 s after onset.

Relative to the reach-free comparison, this decline was evident at the session level (−0.406 ± 0.094 versus +0.041 ± 0.019 z; W = 79, P = 3.8 × 10⁻⁵) but did not reach significance when mice were the unit of analysis (−0.396 ± 0.167 versus +0.042 ± 0.024 z; W = 2, P = 0.094), despite the same direction of effect in five of six mice. The reach-free average showed no comparable temporal profile, supporting an association between the ACh signal and reaching. Because animals might move more in the period before a reach, we repeated the comparison using reach-free windows matched for motion energy, a frame-to-frame measure of gross movement from the behavioral video. The elevation remained after matching (+0.442 ± 0.115 versus −0.085 ± 0.037 z; paired difference, +0.527 ± 0.124 z; W = 0, P = 0.031; n = 6 mice; **Figure 4D**). Matching reduced the elevation by 8%, and the residual imbalance in motion energy was 0.17%.

We next asked whether either component of the signal depended on the outcome of the reach. The pre-onset elevation and the subsequent decline were both present in all five outcome categories, including attempts in which no pellet was consumed (decline shown, **Figure 4E**). Mouse-level analysis detected no difference among categories in the elevation (Friedman χ² = 6.40, P = 0.17) or the decline (χ² = 3.04, P = 0.55; n = 5 mice with all categories represented), nor between successful and unsuccessful reaches (elevation, W = 6, P = 0.44; decline, W = 4, P = 0.22; n = 6). A comparison of the full peri-reach signal between successful and unsuccessful attempts detected no difference at any latency (cluster-mass permutation test, P = 0.215). Thus, neither component required successful retrieval or pellet consumption, although these comparisons do not exclude contributions from reward expectation or other events associated with reaching.

We also quantified gross movement directly. Motion energy, measured across 696 reaches in 33 sessions, was lower before reaches than during reach-free periods in all six mice (4.47 ± 0.51 versus 5.02 ± 0.74 arbitrary units; mean within-animal ratio, 0.909 ± 0.026; W = 0, P = 0.031; **Figure 4F**). Gross movement therefore changed in the direction opposite to that required to explain the pre-onset elevation, and the measured movement did not account for it.

Finally, we examined whether the reach-aligned signal changed with training (**Figure 4G**). The profile was evident during both early and late training, with no detectable difference in pre-onset amplitude (+0.640 ± 0.252 versus +0.437 ± 0.070 z; Mann–Whitney U = 154, P = 0.59) or the subsequent decline (−0.448 ± 0.189 versus −0.383 ± 0.107 z; U = 149, P = 0.72). A paired mouse-level comparison likewise detected no change in the decline (W = 9, P = 0.84). The peak of the early-training average appeared higher, but early sessions contained fewer reaches than late sessions (median 4 versus 24), making peak estimates more sensitive to sampling variability. Subsampling sessions to four reaches reduced the early to late peak difference from 0.358 to 0.088 z (P = 0.88). Complementary measures of signal timing also detected no training-related change, including peak latency (−0.72 versus −0.80 s, P = 0.69), the centroid of the positive excursion (−0.32 versus −0.57 s, P = 0.44), and the first crossing of half-maximum (−1.50 versus −1.49 s, P = 0.69). However, only 71 reaches were recorded during early training compared with 629 during late training, limiting our ability to resolve changes across training.

These results identify a reach-associated ACh elevation before paw movement, followed by a decline during execution. The elevation persisted after matching for gross movement, and the decline was observed in both successful and unsuccessful attempts. Although movement trajectories and speed profiles changed with practice, we detected no corresponding change in the reach-aligned ACh profile. This establishes the temporal relationship between cholinergic signaling and reaching. We therefore tested whether CIN inhibition affects movement refinement and the influence of recent outcomes on subsequent reaches.

We expressed the inhibitory DREADD hM4Di in striatal CINs using the Chat-Cre mouse line and administered deschloroclozapine (DCZ; 50μg/kg) during training on the single-pellet task. Control mice expressed mCherry alone in CINs and received the same DCZ treatment (hM4Di, n = 35; control, n = 23)(**Figure S2A**). We examined both the refinement of movements across training and the influence of recent outcomes on subsequent attempts.

Both groups showed movement refinement with practice. Paw speed decreased over portions of the later movement in controls and hM4Di mice (**Figure 5A,B**; cluster-mass permutation tests, control P = 0.007–0.036; hM4Di P = 0.017 and P < 0.0001). Peak speed fell by 9.6% in controls, from 276 to 250 mm/s (P = 0.0024), and by 16.2% in hM4Di mice, from 275 to 231 mm/s (P = 2.6 × 10⁻⁴). Trajectory variability also decreased in both groups, by 16.4% in controls and 24.1% in hM4Di mice (P = 0.011 and P = 4.1 × 10⁻⁶, respectively; **Figure 5C**). The training-related changes in peak speed and trajectory variability did not differ between groups (P = 0.71 and P = 0.40, respectively). Group comparisons of changes in conditional performance likewise detected no difference in aim, grasp, or withdrawal rates after Holm correction (P = 0.23, P = 1.00, and P = 1.00), and comparisons of the full speed and variability profiles pooled across outcomes were not significant. Thus, CIN inhibition did not prevent the principal forms of movement refinement identified in these analyses.

**Figure 5.**
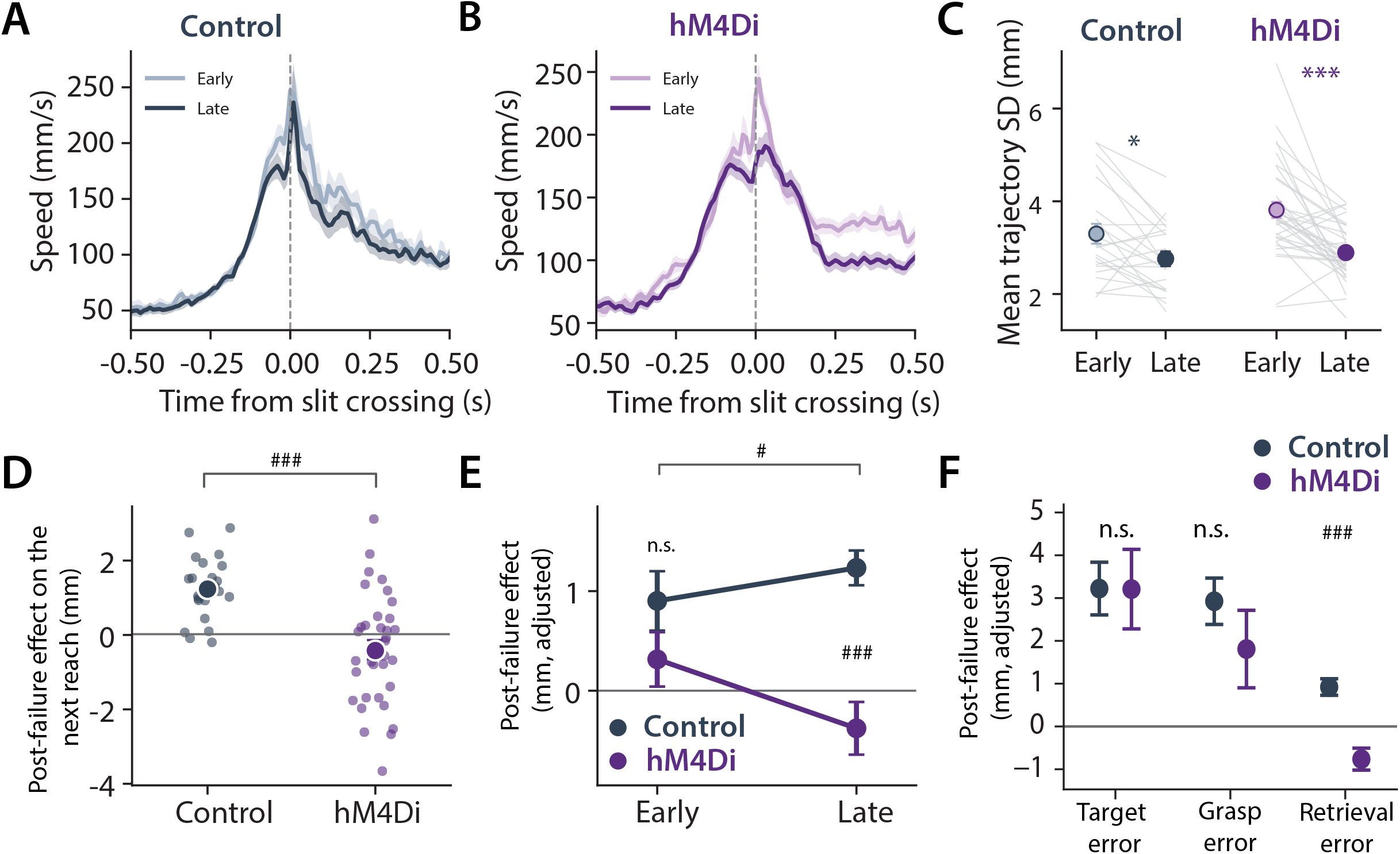
CIN inhibition preserves substantial movement refinement but alters outcome-dependent changes in subsequent reaches. **(A,B)** Paw speed aligned to front-panel crossing (zero) in control mice (grey) and mice expressing hM4Di in striatal CINs (purple), pooled across the five outcome categories. Light and dark shades indicate early (days 1–2) and late (days 9–10) training. Traces show mean ± s.e.m. across mice. Grey bands indicate significant early–late differences identified using pointwise paired t-statistics and cluster-mass permutation tests with 5,000 sign-flips. Control clusters occurred at −0.07 to 0.00 (P = 0.007), +0.03 to +0.08 (P = 0.016), and +0.12 to +0.17 (P = 0.036). hM4Di clusters occurred at −0.03 to +0.04 (P = 0.017) and +0.20 to +0.50 (P < 0.0001). Intervals are expressed in seconds relative to front-panel crossing. **(C)** Trajectory variability during early and late training, calculated as the root-sum-square of across-trial positional standard deviations in the forward and vertical coordinates and averaged over the trajectory. Grey lines connect values from individual mice. Variability decreased from 3.30 to 2.76 mm in controls (−16.4%, P = 0.011) and from 3.81 to 2.89 mm in hM4Di mice (−24.1%, P = 4.1 × 10⁻⁶). The early–late change did not differ significantly between groups (P = 0.40). (**D**) Adjusted effect of a failed attempt on the next reach during late training. For each mouse, the distance of the next reach from its mean successful trajectory was compared following failed and successful attempts, accounting for the preceding reach’s distance, the next reach’s outcome category, and the log inter-trial interval. Positive values indicate greater deviation from the successful trajectory following failure. The effect was +1.233 ± 0.174 mm in controls (21 of 23 mice positive, P < 10⁻⁴) and −0.419 ± 0.260 mm in hM4Di mice (15 of 34 positive, P = 0.13). The groups differed significantly (P = 4.6 × 10⁻⁵). **(E)** Outcome-dependent effect defined in (D), separated by training phase for mice contributing both early and late values (n = 56). The group difference was not significant early (P = 0.18) and was significant late (P = 4.6 × 10⁻⁵); group-by-training-stage interaction, P = 0.034. **(F)** Adjusted effect on the next reach separated by the preceding failure category, relative to a first-attempt success. Effects following target, grasp, and retrieval errors were +3.22, +2.93, and +0.92 mm in controls and +3.21, +1.81, and −0.76 mm in hM4Di mice. Between-group comparisons gave Holm-adjusted P values of 0.46, 0.16, and 1.0 × 10⁻⁴, respectively. The per-mouse linear contrast across successive failure stages was −1.150 ± 0.290 mm per stage in controls (P = 5.5 × 10⁻⁴, n = 23) and −1.986 ± 0.404 mm per stage in hM4Di mice (P = 1.4 × 10⁻⁵, n = 31); these contrasts did not differ significantly between groups (P = 0.278). Control mice expressed mCherry alone; both groups received DCZ during training. Control, n = 23 mice; hM4Di, n = 35. Numbers contributing to each analysis are given per panel. Summary values are mean ± s.e.m., and the mouse was the unit of analysis throughout. Within-group comparisons used two-sided Wilcoxon signed-rank tests, and between-group comparisons used two-sided Mann–Whitney tests. Holm–Bonferroni correction was applied across the three failure-category comparisons in (F). Trial-sequence analyses included 10,124 tracked reaches and 9,892 consecutive pairs. *P < 0.05, ***P < 0.001.

Analysis by outcome category identified a more localized difference in the change in speed during withdrawal (**Figure S2B, C**). For target and retrieval errors, hM4Di mice showed a greater early-to-late reduction in speed than controls over intervals late in the reach (P = 0.0006 and P = 0.005; Holm-adjusted P = 0.006 and P = 0.045 across ten category-by-measure comparisons). However, withdrawal speed was already higher in hM4Di mice during early training, and the groups reached similar values by late training. Accounting for early withdrawal speed detected no group difference in the late-training value (P = 0.168). The larger reduction therefore reflected convergence from different starting points, rather than the acquisition of a slower withdrawal in hM4Di mice. Because DCZ was administered from the first training day, the early difference cannot be separated into an effect of inhibition and a difference between cohorts.

We next asked whether CIN inhibition altered how the outcome of one attempt influenced the movement that followed it. For each mouse we formed all pairs of consecutive attempts and measured how far the second reach of each pair departed from that animal’s mean successful trajectory. These distances were regressed on whether the first reach of the pair had failed, so that reaches following a success served as the reference condition. Because attempts following failures and successes differed systematically in other respects, the model also included the preceding reach’s own distance from the successful trajectory, the outcome category of the second reach, and the interval between the two attempts. The value plotted for each mouse is the resulting coefficient: the difference in deviation between reaches that followed a failure and reaches that followed a success, at matched values of these covariates. A positive value indicates that the next reach deviated further from the successful trajectory following a failure, a negative value that it lay closer to that trajectory, and zero that the preceding outcome had no influence.

During late training, this effect was positive in controls (+1.233 ± 0.174 mm; 21 of 23 mice positive; P < 10⁻⁴), but did not differ significantly from zero in hM4Di mice (−0.419 ± 0.260 mm; 15 of 34 mice positive; P = 0.13). The groups differed significantly (-1.65mm; P = 4.6 × 10⁻⁵; **Figure 5D**). This difference was not detectable during early training and was larger during late training, as supported by a group-by-training-stage interaction (P = 0.034; **Figure 5E**). The late-training group difference persisted when the successful-trajectory reference was constructed from early-training trials only (−1.06 mm, P = 0.002), when the earlier hM4Di sub-cohort was excluded (−1.59 mm, P = 9.4 × 10⁻⁵), and when multiple-attempt next reaches were included (−1.53 mm, P = 9.2 × 10⁻⁵).

Trajectory variability during late training was similar between groups (2.76 versus 2.89 mm; P = 0.35; **Figure 5C**). This suggests that the group difference reflects the influence of the preceding outcome rather than a change in general trial-to-trial variability.

Separating failures by the stage at which they occurred revealed that the group difference was largest after retrieval errors (**Figure 5F**). In controls, the next reach deviated from the successful trajectory following target, grasp, and retrieval errors by +3.22, +2.93, and +0.92 mm, respectively. The corresponding effects in hM4Di mice were +3.21, +1.81, and −0.76 mm. Group differences were not significant after target or grasp errors (Holm-adjusted P = 0.46 and P = 0.16) but were significant after retrieval errors (Holm-adjusted P = 1.0 × 10⁻⁴). Both groups retained a relationship between the stage of failure and the subsequent movement, with the largest deviations following earlier failures. The per-mouse linear trend across failure stages was significant in controls (−1.150 ± 0.290 mm per stage; P = 5.5 × 10⁻⁴) and hM4Di mice (−1.986 ± 0.404 mm per stage; P = 1.4 × 10⁻⁵) and did not differ significantly between groups (P = 0.278). CIN inhibition therefore did not abolish sensitivity to the stage of failure but altered the subsequent movement following attempts in which the pellet had been secured and then lost.

Together, these findings distinguish refinement of the practiced movement from the influence of recent outcomes on subsequent attempts. Mice undergoing CIN inhibition developed more consistent trajectories and showed substantial changes in movement speed across training. Nevertheless, the effect of failed retrieval on the following reach differed from controls. These results support a contribution of striatal CINs to outcome-dependent changes in movement. Because inhibition was applied throughout training, they do not distinguish an acute effect on the use of outcome information from an effect on how that relationship developed with practice.

## Discussion

In this study we resolved skilled reaching into the stages at which individual attempts succeeded or failed and related these outcomes to changes in movement kinematics and striatal cholinergic signaling. Practice improved the probability of contacting the pellet and withdrawing it to the mouth, while successful grasp after contact remained relatively stable. Paw trajectories became more reproducible across several successful and unsuccessful outcomes, whereas changes in the mean speed profile were detected in successful reaches. Striatal ACh was elevated before reach onset and declined during execution in both early and late training. Chemogenetic inhibition of CINs preserved movement refinement but altered the influence of recent outcomes on the next attempt, most clearly after the pellet had been grasped and then lost. These findings identify separable changes in the execution of a practiced movement and its adjustment following an unsuccessful attempt and implicate local cholinergic signaling in the latter.

Skilled reaching requires coordination of forelimb transport, paw orientation, digit closure and withdrawal of the secured object. Detailed descriptions of rat reaching have long recognized that these movements form an ordered sequence whose individual components can be disrupted independently (Whishaw and Pellis, 1990). Their coordination does not require that each component improve at the same rate. During reach-to-grasp learning, gross forelimb measures become consistent before fine movements have completed their refinement, and these changes have partially distinct neural correlates and requirements (Lemke et al., 2019; Bova et al., 2021). Our finding that targeting and withdrawal improved while conditional grasp success remained relatively stable further supports the idea that the components of skilled reaching undergo different degrees of refinement with practice.

The sequential organization of reaching also changes how errors should be interpreted. A grasp failure can occur only after the paw contacts the pellet, and a retrieval failure only after the pellet has been secured. Consequently, the frequency of a later error among all attempts depends both on performance at that stage and on how often the animal reaches it. This dependence accounted for two otherwise misleading features of our data. Grasp errors became more frequent because improved targeting created more opportunities to fail at grasping, while the overall frequency of retrieval errors remained stable because increased progression through preceding stages offset improved withdrawal. Evaluating each stage only after successful completion of the preceding stage exposed these changes without discarding the relationship to overall success (**Figure 1**). This approach complements detailed movement scoring by distinguishing a change in the probability of completing a component from a change in the number of opportunities to perform it.

Task success and movement consistency also provide different information about learning. Rodents can converge on reproducible reaching trajectories even when their success rate fails to improve, demonstrating that an action can become consistent without becoming effective (Bova et al., 2021). Here, reductions in trajectory variability were evident within target errors and grasp errors as well as first-attempt successes (**Figure 2**). Thus, refinement was not confined to attempts that obtained the reward. The outcome-resolved analysis establishes that the reduction in variability was present within several categories and cannot be explained solely by an increased proportion of successful reaches. At the same time, a consistent unsuccessful reach need not represent progress toward a successful one.

We also found that the temporal refinement of reaching was more selective. First-attempt successes attained peak speed earlier during the approach and subsequently moved more slowly after aperture crossing, without a detectable change in peak speed itself (**Figure 3**). This pattern indicates a redistribution of speed across the movement, rather than a uniform increase or decrease in vigor. Precise reaching depends on regulating speed at particular phases, and cerebellar output can adjust reach velocity and endpoint position according to when it is engaged (Becker and Person, 2019). Direction and speed can also be refined through distinct thalamic circuits during learning of reaches to spatial targets (Sibener et al., 2025). Our data are consistent with spatial reproducibility and temporal organization undergoing different forms of refinement within the same action. In first-attempt successes both changes occurred after aperture crossing: these reaches became slower and more spatially consistent over the same portion of the movement.

These behavioral distinctions are relevant to how corticostriatal circuits support skilled movement. Circuit requirements depend on the action and its training history. Motor cortex can become dispensable for execution of an extensively trained lever-pressing skill, while dorsolateral striatum remains necessary for its learned kinematics (Kawai et al., 2015; Dhawale et al., 2021). Dexterous reaching retains quite some cortical dependence, and striatal perturbations can alter both forelimb trajectories and target accuracy (Lemke et al., 2019; Lopez-Huerta et al., 2021). The striatum therefore participates in specifying detailed movements as well as learning them.

ACh is particularly well positioned to regulate how incoming information influences these striatal circuits because it is released primarily by CINs within the striatum. Cholinergic signaling regulates glutamatergic transmission, SPN responsiveness and synaptic plasticity, providing a means for afferent activity to modify processing within the same circuit (Ding et al., 2010; Abudukeyoumu et al., 2019). Corticostriatal potentiation can depend on the coincidence of a CIN pause, dopamine activation and SPN depolarization (Reynolds et al., 2022). These mechanisms provide a rationale for examining ACh around individual reaches, while leaving open whether its principal contribution is to movement execution, learning or the use of outcome information.

The pre-reach ACh elevation we observed suggests that cholinergic signaling may influence motor execution before the overt action begins. Coordinated CIN activity occurs around transitions between spontaneous movement states, and direct measurements of ACh reveal ongoing fluctuations organized by glutamatergic afferents even outside explicitly rewarded events (Howe et al., 2019; Krok et al., 2023). In our task, the elevation preceded reach onset despite reduced gross movement during this period and these distinct ACh dynamics were not present during periods of the task where the mouse did not make a forelimb reach but had similar motion energy (**Figure 4**). Differences in measured gross movement therefore do not appear to account for the elevation. The subsequent decline was observed across successful and unsuccessful outcomes, indicating that pellet consumption was not required for the development of this profile. However, these findings do not exclude reward expectation, task engagement or preparation for reaching. Therefore, these recordings establish the timing of striatal ACh relative to reaching but not its behavioral function.

CIN inhibition did not produce the broad impairment of movement refinement that might be expected if this signal were required for all aspects of skill acquisition. Both groups developed more consistent trajectories and altered their speed profiles, and the changes in conditional performance did not distinguish the groups (**Figure 5**). Basal forebrain cholinergic lesions impair reaching acquisition and associated cortical plasticity in rats, whereas mouse studies have found different requirements for cortical cholinergic input across reaching and rotarod learning (Conner et al., 2003; Li and Hollis, 2021). These findings do not predict that suppressing local striatal CINs will reproduce the consequences of removing cholinergic input to cortex. In fact, studies targeting striatal CINs directly report a comparably selective pattern. Ablation of striatal cholinergic neurons impairs reward-related procedural learning and working memory while leaving rotarod motor learning intact (Kitabatake et al., 2003). Our results support a more specific contribution of striatal CINs under these training conditions, without establishing that striatal ACh is dispensable for motor learning.

The clearest behavioral difference after CIN inhibition concerned how an unsuccessful attempt influenced the next reach. In trained controls, failure was followed by increased deviation from the animal’s own practiced successful trajectory. This relationship was altered in the hM4Di group and the group difference emerged with training (**Figure 5**). Outcome-dependent changes in movement are well established in reinforcement-based motor learning. In human reaching, recent reward history regulates trial-to-trial changes, with larger changes after unrewarded attempts and impaired modulation in Parkinson’s disease (Pekny et al., 2015). Exploration of the workspace also influences the reaching strategies subsequently acquired by mice (Mosberger et al., 2024). Induced reach errors likewise shape subsequent attempts, where closed-loop perturbation of a specific cerebellar output pathway produced learned shifts in the timing of future submovements (Dobrott et al., 2026). The cerebellum influences striatal activity through a disynaptic pathway (Hoshi et al., 2005; Chen et al., 2014), and thalamic input is the principal excitatory drive onto striatal CINs (Lapper and Bolam, 1992; Ding et al., 2010), but whether cerebellar signals reach CINs by this route has not been tested. Our findings connect such outcome-dependent adjustments to striatal CIN function within an already practiced action. The measured deviation does not establish that the next reach was better or that the change constituted an adaptive correction. It nevertheless shows that recent experience influenced execution, beyond the gradual refinement observed across training.

The outcome decomposition was important here as well. The group difference was clearest after retrieval errors, when the pellet had been secured but was lost before consumption. Responses after target errors were preserved, and both groups retained a gradient across failure stages. The manipulation therefore did not eliminate all sensitivity to failure or abolish the distinction between early and late failure. Instead, it changed the influence of a particular outcome on the next movement. Because retrieval errors followed successful targeting and grasping, they combined substantial completion of the action with failure to obtain the reward. One possibility is that evaluating such an outcome requires integration of information about the completed movement with the absence of its expected consequence. CINs could contribute to this evaluation, allowing a failure late in the sequence to modify the next reach even when much of the preceding action had been executed successfully. Thus, our data support a contribution to outcome-dependent updating, while leaving the information being evaluated unresolved.

This interpretation extends established roles for striatal cholinergic signaling in behavioral flexibility. Thalamic regulation of CINs in dorsomedial striatum is required when new action-outcome associations must be learned in the presence of existing associations, and putative CIN activity can reflect inferred task states through its dependence on orbitofrontal input (Bradfield et al., 2013; Stalnaker et al., 2016). More recently, spatially resolved ACh measurements linked unexpected non-reward to subsequent switching, with CIN inhibition reducing this response (Sarpong et al., 2025). Recent work additionally links dorsomedial striatal ACh to the construction of new latent states during reversal learning, particularly when non-reward is ambiguous (Lowes et al., 2026). These tasks require changes in associations or choices. In single-pellet reaching, the task contingency remains fixed and the animal repeats the same action. Our findings suggest that cholinergic contributions to the evaluation of outcomes can also operate at the level of updating how that action is executed.

The timing of the ACh elevation raises a specific possibility for this updating mechanism. The relevant signal would be the elevation before the next reach, after the outcome of the preceding attempt is available. On this interpretation, pre-reach ACh reflects the evaluation of recent outcomes to update execution of the upcoming movement. Comparing the cholinergic profile across the outcomes of the current reach does not address this, because it does not condition on the preceding outcome. We therefore compared consecutive attempts directly, restricting the analysis to pairs separated by at least 2 s. Pre-reach ACh did not differ according to whether the previous attempt succeeded or failed (−0.009 z; P = 0.81; n = 5 mice; one mouse contributed a single qualifying attempt following a success and was excluded). Values by stage of the preceding failure were +0.006, −0.087 and +0.227 z after target, grasp and retrieval errors respectively (all P ≥ 0.062). These recordings therefore do not establish that the elevation carries outcome information about the preceding attempt or the current one. ACh could instead regulate the responsiveness of SPNs to outcome-related inputs arriving from elsewhere, allowing the same broadly timed cholinergic signal to have different consequences depending on recent experience. Dopamine is another plausible component of this process, given its outcome-related dynamics during skilled reaching and the progressive effects of precisely timed dopamine manipulations on reach kinematics (Phillips et al., 2024). Distinguishing these possibilities will require timing perturbation of cholinergic activity precisely between successive attempts.

Several limitations are present in our approach. Our paw tracking was restricted to the sagittal plane and did not resolve digit movements. The bulk ACh recordings average signals that can be spatially heterogeneous. Similarly, chemogenetic inhibition throughout training cannot distinguish an immediate requirement for using outcome information from a requirement for acquiring that relationship through training.

However, our study does demonstrate that improvement in skilled reaching comprises distinct changes in the probability of completing successive stages, the reproducibility of paw trajectories and the temporal organization of movement. Separating these features from the influence of recent outcomes revealed a specific contribution of striatal CINs to how a practiced reach is adjusted after an attempt in which the pellet was grasped and then lost. Together with the pre-reach ACh elevation, these findings support a role for local cholinergic signaling in the outcome-dependent adjustment of a learned movement and identify a window before movement onset in which that influence could be exerted.

## Methods

### Animals

Experiments were performed in adult male and female mice aged 4–8 months. Wild-type mice were C57BL/6 (Jackson laboratory). For chemogenetic experiments, ChAT-Cre mice (Jackson Laboratory, stock no. 031661) were crossed with Ai14 reporter mice (B6.Cg-Gt(ROSA)26Sortm14(CAG-tdTomato)Hze/J; Jackson Laboratory, stock no. 007914). Mice used for acetylcholine imaging were wild-type C57BL/6 (Jackson laboratory). Mice were group housed before food restriction and individually housed during food restriction to control daily food intake. Animals were maintained on a 12-h light/dark cycle. Water and food was available ad libitum except during the food-restriction protocol described below. All procedures were approved by the Northwestern Institutional animal care and use committee and conducted in accordance with animal care guidelines.

### Single-pellet reaching task

Mice underwent two to four days of food restriction, two days of habituation, and two days of pre-training before completing ten consecutive days of single-pellet reaching training. Baseline body weight was measured before food restriction started. Male mice weighed approximately 25–40 g and female mice approximately 20–35 g. During food restriction, mice were individually housed to allow precise control of daily food intake. Standard laboratory chow was provided in the home cage. Food intake was adjusted to maintain mice at approximately 90% of baseline body weight during pre-training and above 85% throughout training. During habituation, reward pellets were provided in the home cage to familiarize mice with Bio-Serv 20-mg chocolate- flavored pellets (F0530120; Bio-Serv) as food reward. The reaching apparatus consisted of a chamber with a 0.5-cm-wide vertical aperture and an external 1.2-cm-height shelf on which a pellet was positioned 1 cm from the front wall (**Figure 1A**) The chamber measured 20 cm in depth, 20 cm in height, and 8 cm in width. During pre-training, pellets were positioned progressively farther from the aperture to encourage reaching and grasping. The preferred forelimb was determined from the first ten successful reaches. During training, the pellet was positioned on a small indentation made in the platform at a 45° angle relative to the opening. Mice completed one training session per day. Each session ended when 20 pellets had been successfully retrieved or 10 min had elapsed. Behavioral recording began approximately 10–30 s after the mouse was placed in the chamber. Reward pellets were manually positioned using surgical forceps. Following a successful retrieval, or when a pellet was displaced from its designated position, a new pellet was presented once the mouse was no longer actively reaching through the opening. When a pellet remained in its designated position after an unsuccessful reach, the same pellet remained available for additional attempts. Consequently, mice could make multiple reaching attempts toward the same pellet, and successful retrievals following one or more unsuccessful attempts were retained as multiple-attempt successes. Sessions were conducted 10 am to 8 pm, and at approximately the same time of day for each mouse across the 10 training days. Reaching behavior was recorded in the sagittal plane using high-speed video for outcome scoring and paw tracking.

### Behavioral scoring and reach outcome classification

Individual reach attempts were manually scored from video and assigned to five mutually exclusive outcome categories according to the stage at which the attempt succeeded or failed. Target errors were defined as attempts in which the paw failed to contact the pellet. Grasp errors involved pellet contact without a secure grasp, including attempts in which the pellet was displaced. Retrieval errors occurred when the pellet was grasped but dropped before being returned to the mouth. Successful retrievals were classified as first-attempt successes when the pellet was obtained on the initial reach and multiple-attempt successes when retrieval followed one or more unsuccessful reaches at the same pellet. Target, grasp and retrieval errors are therefore attempts in which the pellet was not obtained, and first-attempt and multiple-attempt successes are attempts in which it was; the cascade formulas below depend on this grouping. For kinematic and behavioral scoring, reach onset is defined as the moment the paw initiates its forward movement toward the aperture, whereas for in vivo calcium and ACh recording analyses, reach start time is specifically identified by a transition frame showing a small paw jitter following a stationary period. A new attempt series (or new pellet trial) is marked when a mouse pauses continuous attempts for more than 2 seconds, often due to shifting attention or walking away, and subsequently returns to resume reaching. Each failed reach preceding a multiple-attempt success was tracked and scored separately as an individual error attempt, with only the final successful reach assigned to the multiple-attempt success category. Furthermore, only successes are tracked regarding whether they follow prior failures or successes, whereas all error trials are pooled together regardless of preceding error history or subsequent success outcomes. Early and late performance were assessed using training days 1–2 and 9–10, respectively. Animals were included irrespective of their change in overall success rate.

Outcome percentages were calculated within each mouse and training phase before averaging across mice. Counts were combined across the two sessions within each phase and the percentage computed from the pooled counts, rather than averaging session-level percentages; per-mouse percentages were then averaged across mice.

### Stage-specific analysis of reaching performance

To assess performance at successive stages independently of changes in the number of attempts progressing to them, outcomes were analyzed as a nested cascade. For each mouse and training phase, aim rate was defined as the proportion of all attempts contacting the pellet, grasp rate as the proportion of pellet-contacting attempts securing it, and withdraw rate as the proportion of successful grasps returning the pellet to the mouth. Thus when N denotes the total number of attempts, and t, g, r, f, and m denote the numbers of target errors, grasp errors, retrieval errors, first-attempt successes, and multiple-attempt successes, respectively, then N = t + g + r + f + m.

Conditional rates were calculated as: Aim rate = (N − t)/N Grasp rate = (f + m + r)/(N − t) Withdraw rate = (f + m)/(f + m + r). The scoring definitions established the sequential structure of this analysis. Thus grasp errors required prior pellet contact, and retrieval errors required a completed grasp. Consequently, the denominator at each successive stage equaled the numerator of the preceding stage, and the product of the three conditional rates equaled overall retrieval success, (f + m)/N, within each mouse and training phase. Rates were calculated within individual mice before group averaging. The mean of the individual products therefore equaled mean overall success.

#### Decomposition of marginal outcome categories

Because the five outcome categories partitioned every attempt, a change in the share of attempts ending in one category arose either because performance at that category’s own stage changed, or because the number of attempts reaching that stage changed. A category’s share of all attempts was the product of the conditional rates leading to it. The grasp-error share equals p_aim x (1 - p_grasp), and the retrieval-error share equals p_aim x p_grasp x (1 - p_withdraw), where p_aim, p_grasp and p_withdraw are the conditional rates defined above. For each mouse, the early-to-late change in a category’s share was decomposed into a component attributable to the preceding stages’ rates and a component attributable to the category’s own stage. Each component was computed by substituting one factor at a time from its early to its late value while holding the other at its early value and then averaged over both orderings of substitution. This is the two-factor Shapley decomposition, and it has the property that neither component depends on the order in which the factors are exchanged and that the two components sum exactly to the observed change in every mouse. The decomposition was performed within each mouse before averaging across mice. Each component was tested against zero using a two-sided Wilcoxon signed-rank test computed by exact enumeration.

Because the three quantities reported per category are not independent, the two components summing to the observed change by construction, they are reported without multiple-comparison correction as a pre-specified decomposition rather than as a family of independent tests. This analysis used 29 mice and 5,215 scored attempts.

### DeepLabCut and trajectory analysis

#### High-speed videography and pose estimation

Forelimb movements were recorded in the sagittal plane using a high-speed camera (The Imaging Source, DMK 33UX273). Videos were recorded at a frame rate ranging from 100 to 180 frames/s with a resolution of 720 × 540 pixels under standard room illumination. Frame timing was taken from the per-frame timestamps written with each recording rather than assumed from the nominal rate, because the realized interval is irregular; in the imaging cohort the nominal rate of 150 frames/s corresponded to a measured mean of 152.1 frames/s with individual intervals ranging from 1 to over 20 ms. Dropped frames therefore appear as gaps in the timestamp series and were handled by binning on elapsed time rather than on frame index. This choice was validated against the imaging clock: mapping by timestamps agreed with the fluorescence time base to 13.1 ms at the median, whereas assuming a fixed frame rate disagreed by 176 ms at the median and by up to 3.4 s. Paw position was tracked using DeepLabCut (v2.2/2.3, TensorFlow backend with a ResNet-50 network architecture). The pose-estimation model was trained on manually labeled frames from 11 videos obtained from different wild-type mice on the first day of single-pellet reaching training. Keypoints comprised the left and right paws/wrists and the nose. Predictions with a likelihood ≥0.9 were accepted. Missing or erroneous tracking points were interpolated. The preferred forelimb identified during pre-training determined which paw trajectory was analyzed for each mouse. Pixel coordinates were converted to millimeters using a linear scaling factor determined from the known physical dimensions of the apparatus (the 0.5-cm aperture width and 1-cm pellet distance from the front wall). Forward position was measured relative to the front panel and vertical position relative to the platform surface.

#### Trajectory alignment and normalization

Individual reaches were aligned to aperture crossing, defined as the point at which the paw crossed the plane of the front-panel aperture. Trajectories were time-normalized and resampled to 101 points spanning the outward reach and withdrawal. Maximum forward extent was identified separately as the greatest forward displacement of the paw and was not used as the alignment landmark. Analyses were performed separately by mouse, session, and outcome category. Session-level values were averaged within each mouse and training phase before group-level comparisons, so the mouse was the unit of analysis.

#### Analysis of trajectory variability

Trial-to-trial trajectory variability was calculated from the standard deviations of forward and vertical paw position across reaches at each normalized trajectory point. The two coordinates were combined as their mean for data in Figure 2C and as the root-sum-square for data in Figure 5C. Sessions containing fewer than three scored reaches in an outcome category were excluded from variability estimates for that category. Whole-trajectory variability was calculated by averaging pointwise variability over the complete normalized movement. Training-related changes were also assessed along the trajectory using pointwise paired t-statistics and cluster-mass permutation tests with 5,000 sign-flips. Points were included in a cluster when |t| exceeded the two-sided critical value at alpha = 0.05 for the available degrees of freedom; cluster mass was the sum of |t| over contiguous suprathreshold points; the null distribution was the maximum cluster mass obtained on each of 5,000 random sign-flips of the per-mouse differences, and the cluster-level P value was the proportion of null maxima greater than or equal to the observed cluster mass. Family-wise error was controlled within each outcome category; clusters were not additionally corrected across the five categories. For between-group comparisons, early-to-late changes were calculated within each mouse and compared using pointwise independent-samples t-statistics and cluster-mass permutation tests with 5,000 permutations of group labels. Holm–Bonferroni correction was applied across the ten outcome-category-by-measure comparisons comprising five categories and two measures, speed and trajectory variability. Correction was applied to the cluster-level P values of the most significant cluster in each of the ten comparisons.

#### Analysis of paw speed and kinematics

Instantaneous speed was calculated from the forward and vertical velocity components using the elapsed time between video frames. Speed profiles were aligned to aperture crossing and resampled onto a common axis of real time relative to crossing, spanning −0.5 to +0.5 s in 101 points at 10-ms intervals.

Six summary metrics were calculated: peak speed, time of peak speed relative to aperture crossing, the integral of speed over +0.1 to +0.3 s after crossing, peak deceleration, and the across-trial standard deviations of peak speed and its timing. The speed integral was expressed in millimeters and represented distance traveled during the specified interval. Mean speed was also assessed over approach, pre-contact, contact, withdrawal, post-withdrawal, and whole-trajectory intervals.

### GRAB-ACh expression and in vivo imaging

#### Viral delivery and GRIN lens implantation

GCaMP8f and GRAB-rACh3.0 were expressed using viral vectors delivered from a silk fibroin coating on the GRIN lens, allowing viral delivery and lens implantation in a single procedure (Jackman et al., 2018). Aqueous silk fibroin (5% w/v) was prepared as previously described. Vectors encoding GCaMP8f (pAAV.Syn.jGCaMP8f.WPRE, 162376-AAV9 from Addgene) and GRAB-rACh3.0 (AAV9-hSyn-rACh3.0, from WZ Biosciences Inc.) under the human synapsin promoter were mixed in a 1:1:2 ratio with the 5% silk fibroin solution. Two 500 nl droplets of the fibroin–virus mixture were applied to the flat surface of a GRIN lens (Inscopix, 1050-005473, 1 mm diameter, 4.0 mm length). The coating was dried at room temperature for at least 1 h and vacuum desiccated overnight at 4 °C. Coated lenses were implanted with the coated surface facing the tissue in left dorsolateral striatum under isoflurane anesthesia. Implantation coordinates were AP +0.10 mm, ML 2.0 mm, and DV 2.4 mm from Bregma. A circular craniotomy of approximately 1.7 mm in diameter was made by drilling four corners and removing the intervening bone. The exposed skull surface was thoroughly dried and scored using a scalpel to enhance adhesive bonding. While maintaining a continuous irrigation of artificial cerebrospinal fluid (ACSF) across the site, the dura mater and overlying cortical tissue were carefully aspirated using a 26-GA blunt needle attached to a vacuum source. Aspiration proceeded downward until the distinctive fibers of the corpus callosum became visible, at which point a finer 30-GA blunt needle was employed to dissect and remove the callosal tract layers until the underlying striatal parenchyma was cleanly exposed. The virus-coated GRIN lens with Inscopix miniscope baseplate was subsequently lowered into the prepared aspiration cavity to a depth of −2.4 mm (measured relative to the brain surface at bregma), aligned and stabilized using a vacuum-connected pipette tip on the stereotaxic frame, and permanently secured to the scored skull using cyanoacrylate and opaque dental cement (Parkell C&B-Metabond). Postoperative analgesia consisted of the administration of meloxicam (Ostilox, NDC 13985-559- 10; 5 mg·kg−1 2x/day for 48 h or daily for a week) and dexamethasone (Bimeda, NDC 61133-089-20; 0.2 mg·kg−1 1x/day for 7 days) via intraperitoneal injection, alongside buprenorphine SR (1 mg·kg−1) and Sulfatrim in the drinking water for 7 days. Mice recovered for at least 21 days before food restriction began, and imaging commenced approximately four weeks after implantation. Prior to food restriction, virus expression and sensor visibility were verified using the Inscopix in vivo imaging system to confirm both GCaMP8f and GRAB-rACh3.0 fluorescence. Food restriction was subsequently initiated, and habituation for the in vivo imaging cohort included an additional handling familiarization protocol to accustom the mice to mounting the Inscopix miniscope, followed by the attachment of a dummy camera head-stage during pre-training to acclimate the animals to the extra weight. Habituation and pre-training were conducted over 4 days, leading directly into the start of the 10-day single-pellet reaching training and imaging sessions approximately four weeks after implantation.

### Imaging acquisition

#### Fluorescence preprocessing

Recordings were processed using custom MATLAB scripts designed to record behavioral video and synchronize frame timing with the Inscopix system acquisition. Interleaved movies were separated into green and red channel streams. Rigid motion correction was estimated from the green channel and applied to both channels because the green fluorescence provided more spatial structure for registration.

A single region of interest spanning the usable imaging field was drawn manually, and mean pixel intensity was extracted to obtain a bulk fluorescence trace for each channel. Raw fluorescence traces were automatically downsampled when original acquisition rates exceeded the target frequency, applying MATLAB’s decimate function for anti-aliasing low-pass filtering.

Baseline fluorescence, F₀, was estimated using a dynamic sliding-window procedure moving minimum followed by Gaussian smoothing. Fluorescence changes were calculated as transient ΔF/F percentage traces ((F_raw − F₀_dynamic) / F₀_dynamic) × 100 and saved for subsequent reach-aligned analysis. For reach-aligned comparisons, fluorescence sequences were interpolated onto a standardized time grid spanning −2.0 to +2.0 s relative to reach onset.

#### Synchronization of imaging and behavior

Reach onset was scored from behavioral video as the first forward movement of the paw.

Video-frame indices were mapped to the fluorescence time base using the acquisition rates and the offset established by the synchronization signal. Actual per-frame timestamps recorded during acquisition were retained and utilized directly for temporal mapping, ensuring that irregular frame intervals or dropped frames were explicitly accounted for without assuming a fixed frame rate.

The mapping was additionally checked by cross-correlating exported per-reach fluorescence sequences with the corresponding whole-session trace and confirming agreement with the expected alignment index when an unambiguous maximum was obtained.

### GRAB-ACh signal analysis

#### Normalization and reach-aligned averages

Bulk ΔF/F₀ traces were z-scored using the mean and standard deviation of each recording session. Whole-session normalization was used because the ACh elevation extended into the pre-reach period, making a local pre-onset baseline unsuitable for quantifying this signal.

For each scored reach, fluorescence was extracted from −2.0 to +2.0 s relative to reach onset. Successful and unsuccessful attempts were additionally compared across the whole peri-reach window using a cluster-mass permutation test on per-animal mean traces, with cluster significance assessed against 5,000 sign-flipped permutations. Original fluorescence traces, acquired at Inscopix system sampling rates and downsampled to a 10 Hz working frequency, were interpolated onto a common 41-point time grid using linear interpolation across the 4-second window. Reach-aligned traces were averaged within each session. Pre-onset fluorescence was quantified as the mean z-score over −2.0 to −0.6 s, and post-onset fluorescence as the mean over +1.0 to +2.0 s. The decline was defined as the post-onset value minus the pre-onset value, with negative values indicating a decrease.

Session-level averages were used to display the fluorescence time course. Mouse-level summaries were also calculated for statistical comparisons. Outcome-resolved plots show mean ± s.e.m. across reaches, whereas comparisons between outcome categories used mouse-level values. Session-level tests were reported separately from mouse-level tests.

#### Reach-free comparison

For each session, surrogate onset times were sampled from periods at least 4 s from any scored reach. Twenty surrogate times per scored reach were drawn uniformly from the eligible recording period. Fluorescence windows were extracted and processed using the same normalization, alignment window, and summary measures as the scored reaches. This produced a reach-free comparison matched to each session’s number of scored reaches. Real and surrogate values were compared within sessions and within mice.

#### Movement analysis and movement-matched comparison

Gross movement was quantified as the mean absolute difference in pixel intensity between successive behavioral video frames within a body region of interest that excluded the pellet, aperture, and door. Motion energy was summarized over the pre-onset window and compared with reach-free windows from the same session.

For the movement-matched analysis, each reach was compared with 20 reach-free windows selected from the same motion-energy decile within its session. Pre-onset fluorescence was summarized for real and matched surrogate windows using the same procedure as the unmatched comparison. These analyses included 696 reaches from 33 sessions with usable synchronized behavioral video.

#### Comparisons across training

Early and late recordings were defined as training days 1–2 and 9–10, respectively. Pre-onset fluorescence and the post-onset decline were compared between phases using session-level values, with paired mouse-level comparisons where reported.

To assess whether unequal numbers of reaches influenced estimates of peak fluorescence, sessions were subsampled to four reaches before calculating reach-aligned averages.

Timing was assessed using the latency of the maximum of the reach-aligned signal, the centroid of its positive excursion, and the first crossing of half-maximum. The lag between early and late averages was estimated by cross-correlation.

### Chemogenetic manipulation of cholinergic interneurons

The inhibitory DREADD hM4Di was expressed in striatal cholinergic interneurons of ChAT-Cre mice using AAV8-EF1a-DIO-hM4D(Gi)-mCherry-WPRE-HGHpA (custom-packaged by Vigene/Vorovice, 2.19 × 10¹³ vg/mL). Control mice received an AAV expressing mCherry alone (pAAV8-Ef1a-mCherry, Addgene plasmid #114470, titer 1 × 10¹³ vg/mL). Bilateral microinjections were targeted to the dorsolateral striatum across multiple sites to ensure broad column coverage (coordinates relative to bregma: [1] lateral 2.30 mm, posterior 1.00 mm; [2] lateral 1.50 mm, posterior 1.00 mm; [3] lateral 1.70 mm, posterior 0.34 mm). For each injection site, the needle was lowered to the deeper DV coordinate (3.40 mm), and infusions were performed by delivering a total volume of 1 µL over 10 minutes while a stereotaxic automated injector simultaneously withdrew the needle upward by 0.6 mm (to a final depth of 2.80 mm) over the same 10-minute duration. Following the infusion, the needle was left in place for a 5-minute dwell time to prevent backflow before complete withdrawal. Surgical anesthesia and postoperative care included the administration of meloxicam (Ostilox, NDC 13985-559-10; 5 mg·kg⁻¹ 2x/day for 48 h or daily for a week) via intraperitoneal injection, alongside buprenorphine SR (1 mg·kg⁻¹). Triple antibiotic ointment was applied topically to the sutured scalp incision, and mice were monitored daily during recovery.

Mice began behavioral training after a 3–4 week recovery and expression period. Both hM4Di and control mice received deschloroclozapine (DCZ) during training. DCZ powder was first dissolved in DMSO and diluted in sterile saline to a final concentration of 2% DMSO in saline. Mice received an intraperitoneal injection of DCZ (50µg·kg⁻¹) or vehicle (2% DMSO in saline) exactly 30 minutes prior to the start of each daily training session.

### Statistical analysis

The mouse was the unit of biological replication for behavioral and kinematic analyses. Early (days 1–2) and late (days 9–10) training were compared within animals unless otherwise indicated. Summary values are reported as mean ± s.e.m. of per-mouse values, except where fluorescence plots summarize sessions or individual reaches, as specified in the figure legends. ACh analyses included mouse-level comparisons and supplementary session-level comparisons; sessions from the same mouse were not treated as independent biological replicates in mouse-level analyses.

Paired comparisons used two-sided Wilcoxon signed-rank tests, and independent-group comparisons used two-sided Mann–Whitney tests. Comparisons among reach-outcome categories within mice used Friedman tests.

Multiple comparisons were corrected using the Holm–Bonferroni procedure within defined families. Separate families comprised the five reach-outcome categories and the three conditional stages of the behavioral cascade. Between-group comparisons of changes in speed and trajectory variability across outcome categories were corrected across ten category-by-measure comparisons. Outcome-history comparisons between groups were corrected across the three preceding failure categories. Uncorrected and adjusted P values are identified in the text and legends. Overall retrieval success was analyzed separately from the conditional stage comparisons.

Where directional consistency across mice was assessed statistically, the number showing improvement was compared with a null proportion of 0.5 using a two-sided exact binomial test. Confidence intervals for mean paired differences were calculated as t-based 95% intervals. Percentile bootstrap intervals were calculated using 10,000 resamples of mice where specified.

Pointwise comparisons of trajectory variability and paw-speed profiles used cluster-mass permutation tests. Within-mouse early–late comparisons used paired t-statistics and 5,000 sign-flips. Between-group comparisons of per-mouse early–late changes used independent-samples t-statistics and 5,000 permutations of group labels.

For each mouse, a reference trajectory was defined as the mean first-attempt-success trajectory across the days analyzed. The distance of an individual reach from this reference was the root mean square of the point-by-point Euclidean deviation, computed across the entire normalized trajectory. Outcome-history effects were estimated while accounting for the preceding reach’s distance from this reference, the next reach’s outcome category, and the log inter-trial interval. Statistical analyses were performed using SciPy 1.17.1 and statsmodels 0.14.6. All tests were two-sided unless otherwise specified, with P < 0.05 used as the significance threshold after correction where applicable.

## Acknowledgements

This work was supported by NIH/NIMH R01MH099114 award to AC. We thank Dr Jones Parker for detailed comments on the manuscript

## Author Contributions

N-HY, JJM, AC designed the experiments N-HY, JJM and LP performed experiments and analyzed data. N-HY, JJM, AC prepared the manuscript.

---

**Figure S1.**
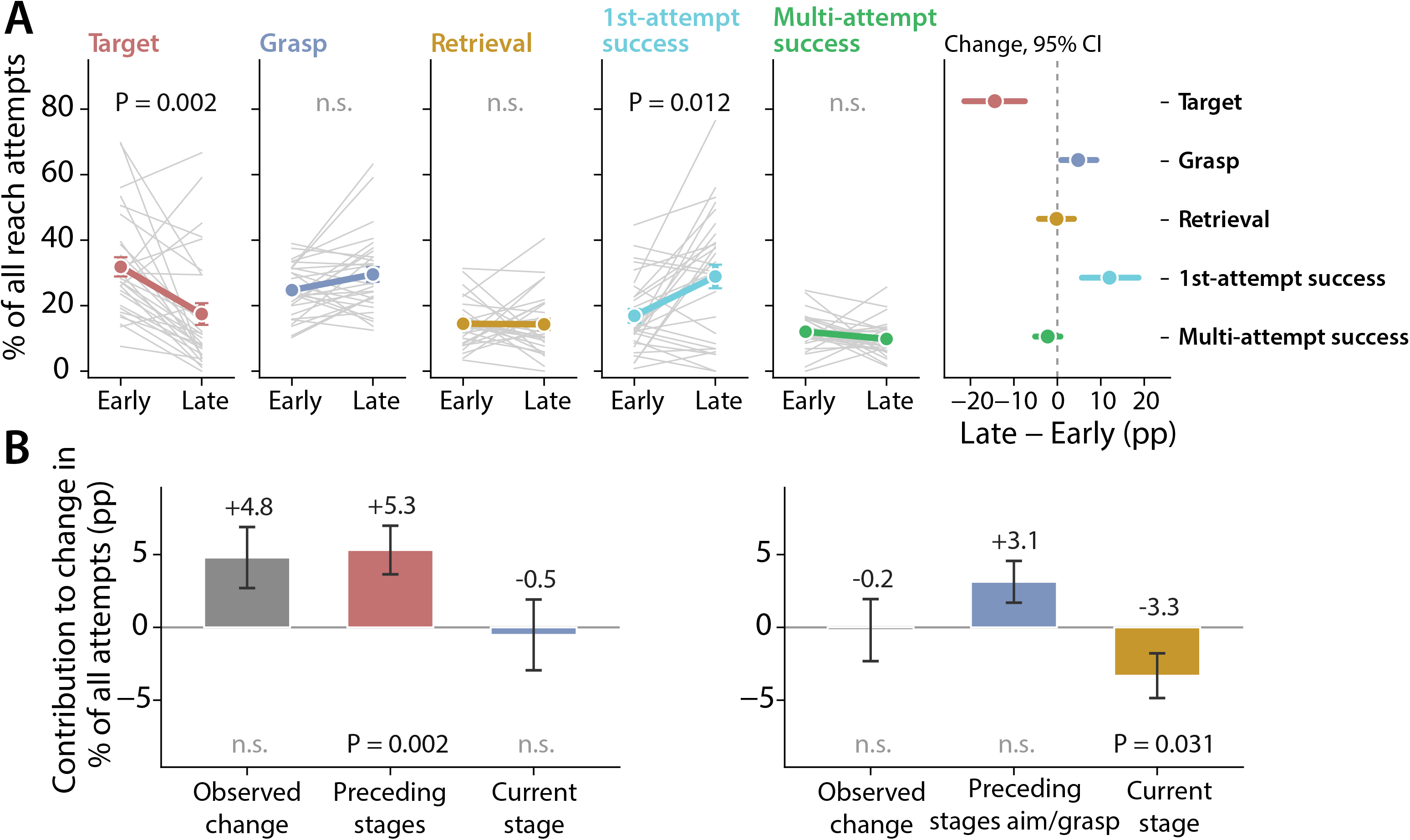
Changes in reach outcomes and contributions from successive stages of the action. **(A)** Percentage of all reach attempts assigned to each outcome category during early (days 1– 2) and late (days 9–10) training. Grey lines connect values from individual mice; colored symbols and lines show mean ± s.e.m. Early versus late comparisons: target errors, W = 61.0, P = 3.7 × 10⁻⁴, P_Holm = 0.0018; grasp errors, W = 128.0, P = 0.053, P_Holm = 0.160; retrieval errors, W = 209.0, P = 0.865, P_Holm = 0.865; first-attempt successes, W = 84.0, P = 0.003, P_Holm = 0.012; multiple-attempt successes, W = 141.0, P = 0.101, P_Holm = 0.202. Plot annotations show Holm-adjusted P values. Right, mean paired change (late minus early) in percentage points, with 95% percentile bootstrap confidence intervals calculated from 10,000 resamples of mice. The dashed line indicates zero change. **(B)** Decomposition of changes in grasp-error and retrieval-error percentages into contributions from performance at preceding stages and performance at the current stage. Components were calculated within each mouse and sum to its observed change. For grasp errors, the preceding-stage contribution reflects changes in aim performance; for retrieval errors, it reflects changes in aim and grasp performance. A negative current-stage contribution indicates improved conditional performance, reducing the proportion of attempts ending in that error. Bars show mean ± s.e.m.; labels give mean changes in percentage points. Grey bars denote observed changes, and colored bars denote the component contributions. For grasp errors, the observed change was +4.8 ± 2.1 percentage points (W = 128.0, P = 0.053), comprising a preceding-stage contribution of +5.3 ± 1.7 (W = 80.0, P = 0.002) and a current-stage contribution of −0.5 ± 2.4 (W = 208.0, P = 0.85). For retrieval errors, the observed change was −0.2 ± 2.1 percentage points (W = 209.0, P = 0.865), comprising a preceding-stage contribution of +3.1 ± 1.4 (W = 128.0, P = 0.053) and a current-stage contribution of −3.3 ± 1.5 (W = 118.0, P = 0.031). Data comprise 5,215 scored attempts from 29 mice. The mouse was the unit of analysis throughout. Two-sided Wilcoxon signed-rank tests compared early and late values in (**A**), with Holm–Bonferroni correction across the five outcome categories. In (**B**), two-sided Wilcoxon signed-rank tests assessed observed changes and component contributions against zero; P values are uncorrected. n.s., P ≥ 0.05 using the applicable corrected or uncorrected value.

**Figure S2.**
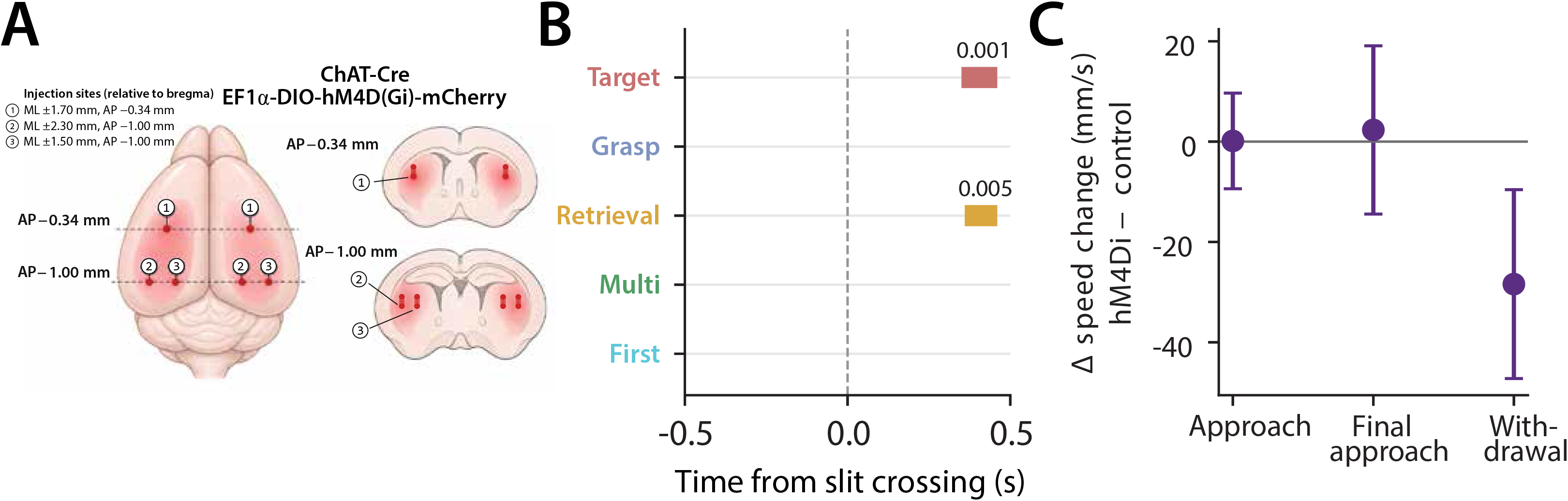
Outcome-resolved kinematic comparisons. (**A**) Schematic illustrating bilateral viral delivery of AAV8-EF1α-DIO-hM4D(Gi)-mCherry into the dorsolateral striatum of ChAT-Cre mice. Dorsal view (left) and coronal sections (right) show the three injection sites in each hemisphere, with coordinates relative to bregma: site 1, AP −0.34 mm, ML ±1.70 mm; site 2, AP −1.00 mm, ML ±2.30 mm; and site 3, AP −1.00 mm, ML ±1.50 mm. For each injection site, the needle was first lowered to a deep coordinate at 3.40 mm (DV), and infusions were performed by delivering a total volume of 1 µL over 10 minutes while the injector needle was simultaneously withdrawn by 0.6 mm to a final depth of 2.80 mm (DV). (**B**) Intervals in which the early-to-late change in paw speed differed between control and hM4Di mice, separated by outcome category. Comparisons used independent-samples t-statistics on per-mouse changes and cluster-mass permutation tests with 5,000 permutations of group labels. Significant clusters occurred during withdrawal for target errors (+0.35 to +0.46, P = 0.0006, P_Holm = 0.006) and retrieval errors (+0.36 to +0.46, P = 0.005, P_Holm = 0.045). Holm–Bonferroni correction was applied across ten comparisons comprising five outcome categories and two measures, speed and trajectory variability. No significant cluster was detected for trajectory variability in any category or for either measure when profiles were pooled across categories. *Speed intervals are expressed in seconds relative to front-panel crossing*. (**C**) Between-group differences in early-to-late changes, pooled across outcome categories and summarized over approach (−0.50 to −0.31), final approach (−0.15 to 0), and withdrawal (+0.35 to +0.46). Values show hM4Di minus control changes, with 95% confidence intervals. Differences in speed change were +0.1, +2.3, and −28.4 mm/s, respectively. Differences in trajectory-variability change were −0.65, −0.28, and −0.54 mm (P = 0.075, 0.47, and 0.22). Mean withdrawal speed was higher in hM4Di mice during early training (128.9 versus 102.0 mm/s, P = 0.004), but did not differ significantly during late training (98.0 versus 99.5 mm/s, P = 0.50). Adjusting late-training speed for the early value also detected no group difference (P = 0.168). The larger reduction in hM4Di mice therefore reflected convergence from different early values to similar late values. Control, n = 23 mice; hM4Di, n = 35. Early training, days 1–2; late training, days 9–10. The mouse was the unit of analysis throughout.

